# Immune Checkpoint Inhibition-related Neuroinflammation Disrupts Cognitive Function

**DOI:** 10.1101/2024.07.01.601087

**Authors:** Onwodi V. Ifejeokwu, An Do, Sanad M. El Khatib, Nhu H. Ho, Angel Zavala, Shivashankar Othy, Munjal M. Acharya

## Abstract

Combinatorial blockade of Cytotoxic T-lymphocyte associated protein 4 (CTLA-4) and Programmed Cell Death Protein 1 (PD-1) significantly improve the progression-free survival of individuals with metastatic cancers, including melanoma. In addition to unleashing anti-tumor immunity, combination immune checkpoint inhibition (ICI) disrupts immune-regulatory networks critical for maintaining homeostasis in various tissues, including the central nervous system (CNS). Although ICI- and cancer-related cognitive impairments (CRCI) in survivors are increasingly becoming evident, our understanding of ICI-induced immune-related adverse effects (IREA) in the CNS remains incomplete. Here, our murine melanoma model reveals that combination ICI impairs hippocampal-dependent learning and memory, as well as memory consolidation processes. Mechanistically, combination ICI disrupted synaptic integrity, and neuronal plasticity, reduced myelin, and further predisposed CNS for exaggerated experimental autoimmune encephalomyelitis. Combination ICI substantially altered both lymphoid and myeloid cells in the CNS. Neurogenesis was unaffected, however, microglial activation persisted for two-months post-ICI, concurrently with cognitive deficits, which parallels clinical observations in survivors. Overall, our results demonstrate that blockade of CTLA-4 and PD-1 alters neuro-immune homeostasis and activates microglia, promoting long-term neurodegeneration and driving cognitive impairments. Therefore, limiting microglial activation is a potential avenue to mitigate CNS IRAE while maintaining the therapeutic benefits of rapidly evolving ICIs and their combinations.

**SIGNIFICANCE:** Despite the superior therapeutic efficacy of immune checkpoint inhibition (ICI) for cancers, its undesired effects on brain function are not fully understood. Here, we demonstrate that combination ICI elevates neuroinflammation, activates microglia, leading to detrimental neurodegenerative and neurocognitive sequelae.

## INTRODUCTION

Immunotherapies have revolutionized cancer treatment and improved overall survival. Immune Checkpoint Inhibition (ICI) therapies, which target T cell negative regulatory pathways have been shown to significantly increase therapeutic response of cancers and improve life expectancy, including primary and metastatic melanoma (Buder-Bakhaya and Hassel, 2018; Hargadon et al., 2018). Commonly used ICI include the blockade of cytotoxic T-lymphocyte associated protein 4 (CTLA-4) and Programmed cell death protein 1 (PD-1) pathways, alone or in combination to rejuvenate exhausted immune cells, including cytotoxic (CD8^+^) T cells that orchestrate tumor cell killing (Buder-Bakhaya and Hassel, 2018; Hargadon et al., 2018). Since the first FDA approval of anti-CTAL-4 (Ipilimumab) in 2011 (Lee et al., 2022), clinical cancer immunotherapies have been evolving, and over half a dozen ICIs have been approved. Despite the success of ICI, off-target effects and normal tissue toxicities, commonly referred to as Immune-Related Adverse Events (IRAE), are reported (Darnell et al., 2020; Das and Johnson, 2019). Although IRAE in the gastrointestinal tract is well characterized (e.g., ICI-associated Colitis), our understanding of IRAE mechanisms in the central nervous system (CNS) is largely incomplete.

Cancer survivors who underwent ICI report headache, fatigue, fever, and loss of appetite (Tarhini, 2013), and emerging clinical data suggest both short-term and long-term adverse effects of ICI on brain functions (Blansfield et al., 2005; Goldberg et al., 2016; Tarhini, 2013). ICI-induced short-term neurotoxicity is commonly reported in patients with melanoma, small cell lung cancer (SCLC), non-small cell lung cancer (NSCLC), and Merkel-cell carcinoma at five weeks post-therapy (Duong et al., 2021). Long-term neurocognitive complications after ICI treatment (11 months up to 2 years) are reported in about 41% of the metastatic melanoma survivors. Longitudinal studies assessing the cognitive function and psychosocial impact of Ipilimumab (anti-CTLA-4) on first-generation survivors revealed numerous psychological issues, including anxiety, depression, and PTSD, as well as impairment in neurocognitive function (Rogiers et al., 2020a; Rogiers et al., 2020b; Rogiers et al., 2023). ICI-related neurotoxicities are often associated with demyelination, CNS autoimmune reaction, and sensorimotor polyneuropathy. Altogether, IRAE following ICI therapy on CNS function is evident. However, the cellular mechanism of how ICI orchestrates these neurodegenerative and neurocognitive complications is unclear (Duong et al., 2021; Fleming et al., 2023; Myers et al., 2023).

A combination of anti-CTLA-4 and anti-PD-1 treatment is highly efficacious for treating melanoma (Curran et al., 2010) and is being tested in several clinical trials for various cancers (Sznol and Melero, 2021; Xiang et al., 2022). Cancer survivors who are receiving this dual ICI (CTLA-4 and PD-1 blockade) are likely to have neurocognitive complications, and we hypothesize that disruption of immune regulatory networks during and post-ICI elevates a proinflammatory environment, including microglial activation, that is detrimental to CNS functions. Microglia, constituting about 12% of all CNS cell types, serve as a CNS resident immune cell. Microglia play both reparative and damaging roles following injury, infections, or neurodegenerative events (Paolicelli et al., 2022). Our past studies have established the detrimental role of microglial activation leading to neuronal and synaptic loss, and cognitive dysfunction following cranial irradiation and chemotherapies (Acharya et al., 2016a; Acharya et al., 2011; Acharya et al., 2016c; Acharya et al., 2015; Allen et al., 2019; Markarian et al., 2021; Montay-Gruel et al., 2019). However, neuroinflammatory and neurodegenerative events post-ICI are scarcely reported. Therefore, it is clear that the neurobiological underpinnings of ICI therapy-related modulation of brain functions need to be determined to develop mitigation strategies. In this study, using the syngeneic melanoma model in immunocompetent mice and a series of cognitive function tests, we demonstrate that blockade of CTLA-4 and PD-1 alters the immune landscape of CNS, fueling neuroinflammation-mediated neurodegeneration thus severely affecting hippocampal-dependent cognitive function.

## RESULTS

### CTLA-4 and PD-1 blockade eliminates melanoma but alters immune cells in the brain

In a primary melanoma survivor model, where no cancer is present in the brain, the most likely source of inflammation leading to neuroinflammation and subsequent brain injury, is peripheral immune cells invading the CNS.

To evaluate the effect of ICI on CNS, we used a clinically relevant cancer survivor model of melanoma: syngeneic mouse model (D4M-UV2 cells) and combination therapy of CTLA-4 and PD-1 blockade in mice. D4M-UV2 is derived from D4M.3.A by ultraviolet (UV) B radiation, carries a mutational load comparable to human Melanomas, is immunogenic, and responds well to ICI (Lo et al., 2021). Because the parental line D4M.3.A was initially derived from male transgenic mice, adult C57Bl6 WT male mice were used in our study. We induced melanoma using bilateral, subcutaneous injection of D4M-UV2 cells and measured tumor growth and response to ICI therapy using calipers **(Fig. 1A)**. 10 days post-tumor induction, tumor-bearing mice were randomly enrolled into ICI-treated (Melanoma-ICI) or isotype control (Melanoma-Veh) groups. ICI groups received two doses of anti-CTLA-4 (clone UC10-4F10-11, BioXcell) and three doses of anti-PD1 (clone 29F.1A12™, BioXcell) every week for three weeks (**Fig. 1A**, Research Design). Age- and gender-matched control groups were injected with matching volumes and quantities of isotype control antibodies (ITC, Armenian hamster IgG, and rat IgGa). Compared to the ITC-

**Figure 1.**
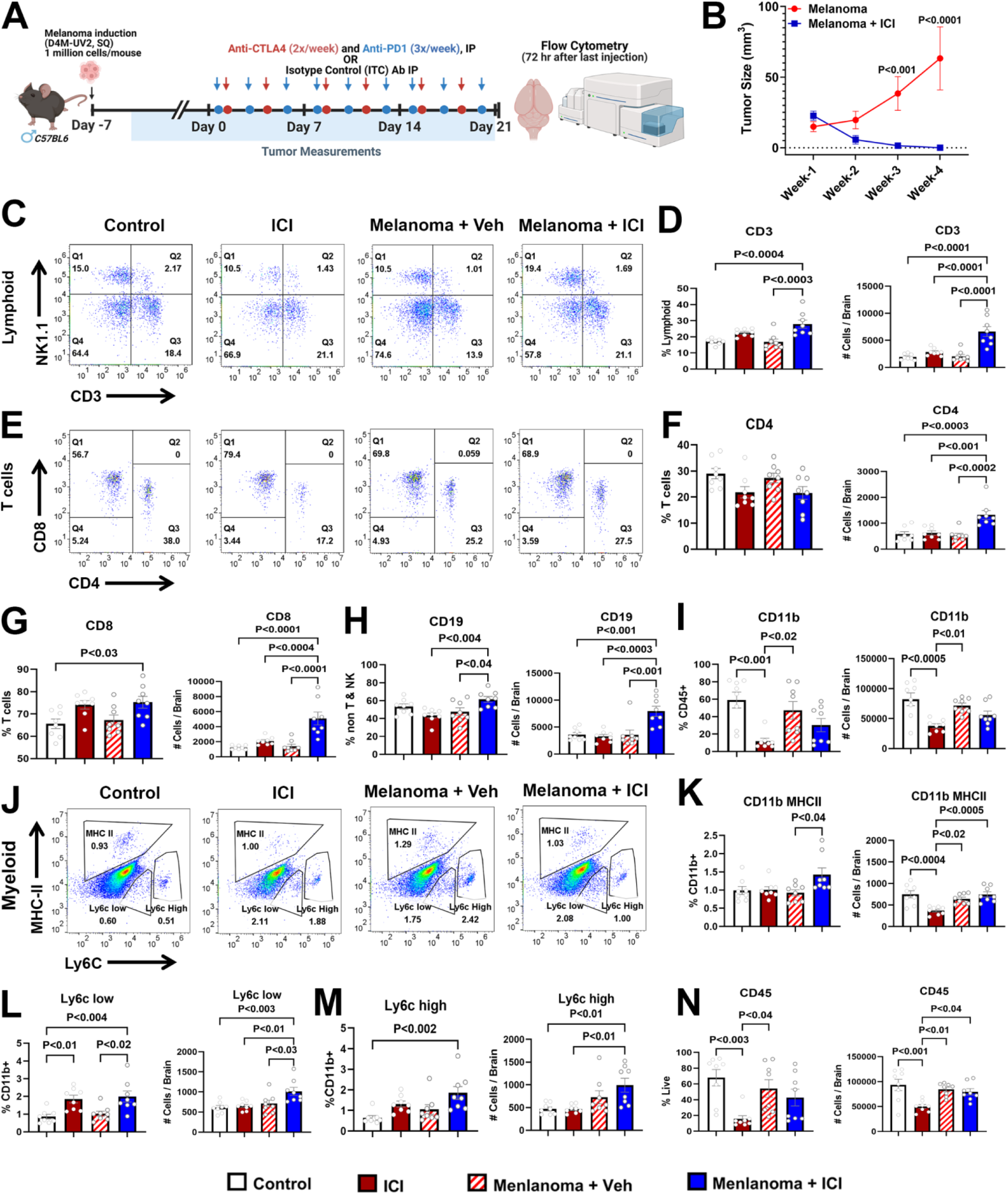
Combination blockade of CTLA-4 and PD-1 eliminates melanoma and induces neuroinflammation. **(A)** Experimental scheme showing study design. C57Bl6 male mice were intradermally injected with 1 million D4M-UV2 cells. Tumors were allowed to engraft for a week before starting ICI treatment and measurements. Melanoma-bearing and control mice were then treated with either vehicle or combinatorial ICI (anti-PD-1 and anti-CTLA4) for 3 weeks. Melanoma tumor measurements (mm^3^) were taken over the course of four weeks throughout ICI treatment. Mice were sacrificed at 72 hours following the last ICI injection to immunophenotype brains using flow cytometry. **(B)** Tumor growth curves showing size of melanomas in control and ICI-treated groups over the course of four weeks. **(C)** Representative dot plots showing NK cells and T cell gating among lymphocytes. **(D)** Frequencies (% of lymphoid) and total number of CD3^+^ T cells from **(C)**. **(E)** Representative dot plots showing CD4 and CD8 T cells among CD3^+^ T cell population. **(F-G)** Frequencies (% of T) and total number of CD4 and CD8 T cells, from **(E). (H-I)** Frequencies and total number of B cells (CD19^+^) and CD11b cells. **(J)** Representative dot plots showing MHCII and Ly6C expression among the myeloid cells. **(K)** Frequencies and number of MHCII^+^ CD11b^+^ cells, from **(J)**. **(L)** Frequencies and number of Ly6c low^+^ cells, from **(J)**. **(M)** Frequencies and number of Ly6c high^+^ cells, from **(J)**. **(N)** Frequencies and number of CD45^+^ cells. Data are presented as mean ± SEM (N=8 mice per group). P values are derived from ANOVA and Tukey’s post hoc test.

treated mice that showed progressive melanoma growth, ICI treatment significantly decreased tumor growth within a week, and by three weeks, tumor mass was undetectable, (*P*’s<0.001, **Fig. 1B**), thus confirming the therapeutic efficacy of dual ICI in our melanoma model.

To determine how ICI broadly changes the immune landscape in the CNS, mice were euthanized 72 hours after the last ICI dosing, and brains were collected after cardiac perfusion **(Fig. 1A)**. We prioritized our analysis of T cells because they are the primary targets of ICI. The gating strategy to quantify the subpopulations of lymphoid and myeloid cells is shown in **Suppl. Fig. S1**. T lymphocyte (CD3^+^) frequencies were nearly 2-fold higher in the Melanoma-ICI group than in non-ICI controls, and the total numbers of T cells were significantly higher in the Mel-ICI group (**Fig. 1C & D**, *P*’s<0.0001). Among T cells, frequencies of helper T (CD4^+^) cells were comparable across groups. However, there was a significant increase in the frequency of cytotoxic T cells (CD8^+^) and the total CD4 and CD8 T cells in the Melanoma-ICI group (**Fig. 1D- G**). We observed a higher frequency and total number of all memory T cells (CD27^+^) in the T cell quadrant (**Suppl. Fig. S2B**). The total number of memory helper T cells was significantly elevated. (**Suppl. Fig. S2D).** Both the frequency and total number of memory cytotoxic T cells were significantly increased in the Melanoma-ICI group. B cell frequency and numbers (CD19^+^, **Fig. 1H, Suppl. Fig. S2G**) and the total numbers of NK cells (CD161^+^) were significantly higher, whereas NK cell frequency was unchanged (**Suppl. Fig. S2H**).

In the myeloid quadrant (CD11b^+^), both patrolling (Ly6C low) and inflammatory (Ly6C high) monocytes significantly increased in the Melanoma-ICI group (**Fig. 1L-M**). Notably, ICI did not result in higher overall CD45^+^ cells (**Fig. 1N**). However, ICI resulted in a higher frequency and number of CD11b^+^ MHCII^+^ cells, suggesting microglial activation. In line with this, frequencies and the number of CD11b^+^ cells were lower, which was reflected in the overall decline of CD45^+^ cells in the ICI group. Together, in consort with the data on the lymphoid cells, these results suggest that combination ICI alters the immune cells in the CNS, primarily T cells and CNS resident myeloid cells (e.g., microglia).

Concurrently, brain (hippocampi) and plasma were collected for a cytokine panel analysis using a seprate cohort of mice (**Suppl. Fig. S3**). We did not find statistical significance in the brain cytokine levels. We found significantly elevated plasma IL1a, TNFa, IL6, IL10, and IL17 in the melanoma + ICI group compared to controls. Plasma IL12 (p70) was elevated in both cancer and non-cancer mice receving ICI treatment.

### Immune checkpoint inhibition impairs cognitive function

Elevated inflammation in the brain (higher memory and cytotoxic T cells and CD11b^+^ MHCII^+^ cells) could culminate into neuronal damage and affect cognitive function. To determine the impact of ICI treatment on learning and memory, memory consolidation, and anxiety-related tasks, a separate cohort of mice underwent a series of cognitive function tests at one-month post-ICI **(Fig. 2A)**. These tests included open field task or activity (OFT), light-dark box test (LDB), object location memory (OLM), and, lastly, fear extinction memory task (FE). EPM and LDB measure levels of anxiety and spontaneous activity. NPR measures hippocampal-dependent episodic memory, and FE measures re-learning and memory consolidation abilities. In the OFT, ICI-treated non-cancer mice exhibited a significant reduction (*P*<0.001) in the percent time spent in the central zone compared to control mice **(Fig. 2B)**. This open field exploration is represented in the heat map of animals from each group exploring the open arenas showing ICI groups spending less time exploring the central zone compared to controls **(Fig. 2C)**. On the LDB test, ICI-treated non-cancer mice exhibited a significant decrease (*P*<0.03) in latency to enter the dark compartment compared to the control mice **(Fig. 2D)**.

**Figure 2.**
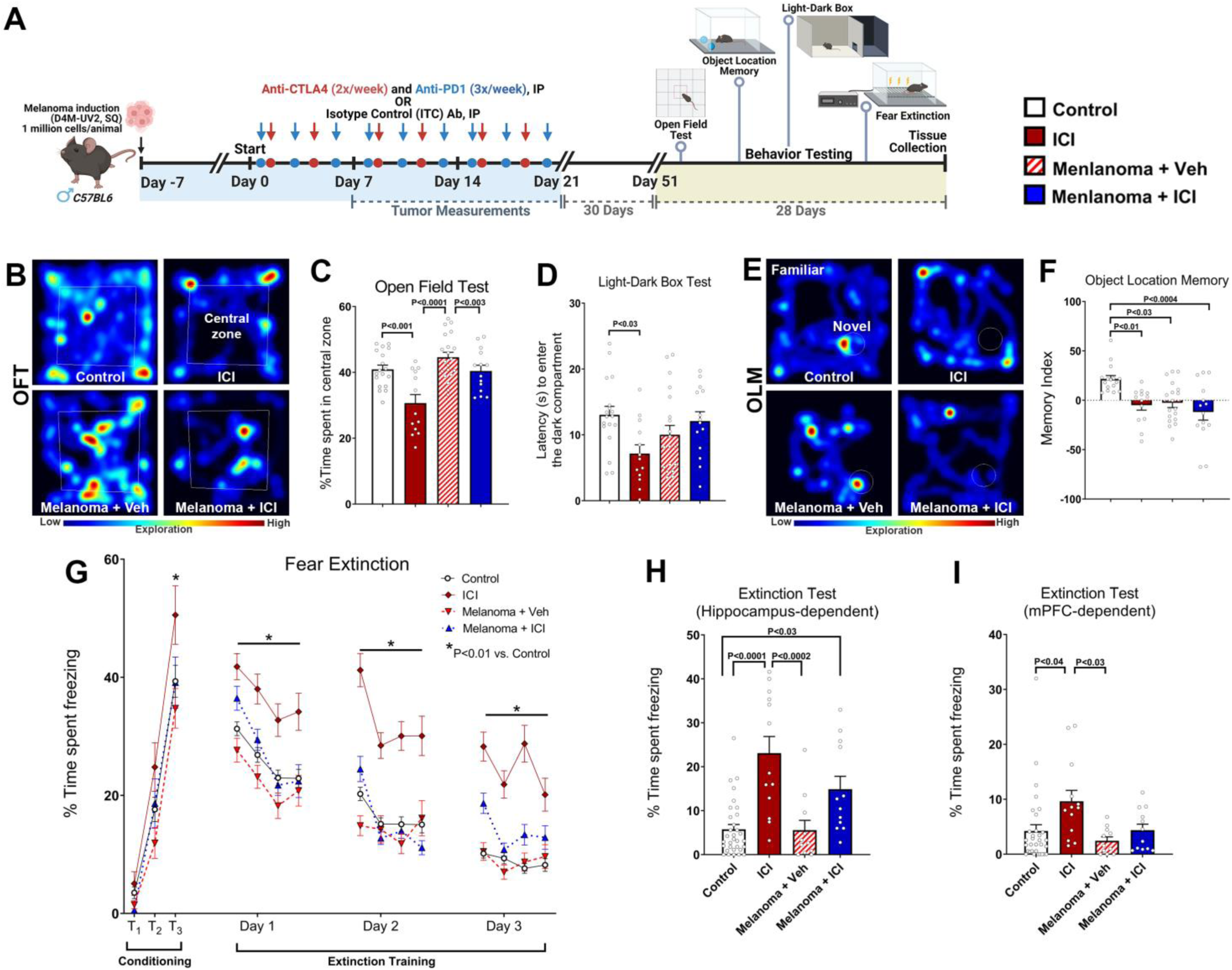
Impaired cognitive function following ICI treatment. **(A)** Research design: Two-month-old C57Bl6 WT male mice received either sham injection or melanoma cells (D4M-UV2, 1 million/animal, SQ). At 7 days post-cancer induction, mice received combinatorial ICI treatment including anti-CTLA4 (twice weekly, IP) and anti-PD-1 (thrice weekly, IP) for three weeks. All mice were administered cognitive function tasks including the Open Field Test (OFT), Light-Dark Box Test (LDB), Object Location Memory test (OLM), and fear extinction memory consolidation test (FE) at 1- month later and euthanized for tissue collection. Paraformaldehyde-fixed brains were cryosectioned to collect 30 μm thick coronal hippocampal sections for immunostaining. **(B-C)** Representative heat maps depicting mice exploring an open field arena during OFT **(B)**. Control mice receiving ICI treatment showed significant reductions in the percentage of time spent in the central zone (35% of the total area) compared to controls treated with a vehicle **(C)**. Melanoma-bearing mice receiving ICI treatment showed reduced central zone exploration compared to melanoma + vehicle mice. **(D)** ICI treatment to the control mice significantly reduced latency to enter the dark compartment, indicating increased anxiety-like behavior, compared to control mice receiving the vehicle. **(E-F)** Representative heat maps depicting mice exploring novel or familiar placement of objects during the OLM task **(E)**. The tendency to explore novel placement of objects was derived from the Memory Index, calculated as: ([Novel object exploration time/Total exploration time] – [Familiar object exploration time/Total exploration time]) ×100. Control mice treated with ICI, melanoma-bearing mice ± ICI treatment showed significantly impaired cognitive function, as indicated by the reduced preference towards the novel placement of objects compared to the control + vehicle group **(F)**. **(G-I)** Cancer-bearing mice ± ICI treatment, and control mice receiving ICI treatment did not impair the acquisition of conditioned fear memory as shown by the elevated freezing following a series of three- tone and shock pairings (80 dB, 0.6 mA, T1–T3, **G**) in a metal grid floor with a mild vinegar odor. ICI- treated control mice showed the highest freezing levels compared to the control + vehicle group (*P<0.01). Subsequently, 24 h later, fear extinction training was administered every 24 h (20 tones) for 3 days in the same environment as during the conditioning day but without administering foot shocks. Each data point for Days 1-3 is presented as the average of percentage of time mice spent freezing for 5 tones (4 data points per day). All mice showed a gradual decrease in freezing behavior (Days 1–3), however, control mice (without cancer) + ICI treatment spent a significantly higher time freezing compared with Control + Vehicle mice. **(H)** Twenty-four hours after the extinction training, on the extinction test, Control + Vehicle and Melanoma + Vehicle mice showed abolished fear memory (reduced freezing) compared with Control + ICI and Melanoma + ICI-treated mice indicating ICI treatment-induced, hippocampal-dependent fear memory consolidation deficit. **(I)** Lastly, at 72 h after the extinction test, mice were tested in a new environment, including a white acrylic floor, additional house light, mild almond odor, and the same tone (60 dB), as on the conditioning day but without foot shock to engage the medial prefrontal cortex (mPFC, **I**). Compared to control + vehicle and melanoma + vehicle groups, control mice (without cancer) receiving ICI treatment showed elevated freezing. Melanoma-bearing mice receiving ICI treatment did not differ in the freezing levels compared to the control + vehicle group. Data are presented as mean ± SEM (*N*=12-32 mice per group). *P* values were derived from two-way ANOVA and Bonferroni’s multiple comparisons test.

To determine the hippocampal-dependent spatial recognition memory, after three days of habituation, animals were administered the OLM task **(Fig. 2E-F)**. This task determines the animal’s ability to explore novel placement of the objects in an unrestricted, non-invasive open environment (Barker et al., 2007; Barker and Warburton, 2011). This activity is calculated as Memory Index [MI = (Novel/Total exploration time) – (Familiar/Total exploration time)] × 100]. A positive MI indicates that animals spent more time exploring novel placement of objects and thus intact hippocampal function. In contrast, zero or a negative MI shows that animals displayed minimal or no preference for the novel object placements and spent equal time exploring familiar and novel places, indicating disrupted hippocampal function. During the test phase, the total time spent exploring familiar and novel objects was comparable for each experimental group. Compared to controls, ICI-treated non-cancer mice (ICI, *P*<0.01), melanoma-bearing mice + ITC (Melanoma + Veh, *P*<0.03), and melanoma-bearing mice treated with ICI (Melanoma + ICI, *P*<0.0004) exhibited significant decreases in the MI. This behavior is represented in the heat map of animals from each group exploring the familiar and novel placement of objects during the OLM test phase showing controls exploring the novel placement higher (red zones) compared to all the other groups **(Fig. 2F)**.

Finally, the mice underwent FE testing. All the groups of mice showed elevated freezing after the initial tone-shock fear conditioning phase (**Fig. 2G**, Conditioning phase, T1-T3). ICI-treated non-cancer mice showed the highest freezing compared to controls (*P*<0.01) after the third tone-shock exposure. At 24h post-conditioning phase, for the next three days, mice were exposed to repeated tones without shock in the same arena with spatial (grid, house light) and odor (vinegar) cues (**Fig. 2G**, Extinction training phase). Mice with intact hippocampal function will exhibit a decrease in freezing behavior during tone as they re-learn from this extinction training by dissociating previously learned freezing response following aversive stimuli (mild foot shocks). ICI-treated non-cancer mice showed elevated freezing compared to controls, indicating a decreased ability to extinguish fear. Finally, during 24-h post-extinction training, an extinction test **(Fig. 2H)** was administered, including exposure to tones in the same spatial and odor-cue environment as the conditioning phase (Day 1). ICI-treated mice showed a significant elevation (*P*<0.0001) in their freezing behavior compared to controls, indicating memory consolidation deficits (**Fig. 2H**). Melanoma + Veh mice showed comparable freezing to control mice, indicating that cancer did not influence fear extinction memory. In contrast, Melanoma + ICI mice showed a significantly elevated freezing (*P*<0.03) compared to the Control + vehicle mice, indicating a detrimental impact of ICI treatment on the fear memory consolidation process. In addition, to test the frontal cortex-dependent extinction memory process, 72 hours after the above extinction test, mice were placed in a box with new flooring (acrylic plate), a spatial cue (extra light), and a new odor (almond). Throughout the 15-minute test, three tones were played without any shock. Again, ICI-treated non-cancer mice exhibited significantly higher freezing behavior than control mice (*P*<0.04). In contrast, Melanoma + Veh and Melanoma + ICI mice did not change the freezing behavior **(Fig. 2I)**. Collectively, these data indicate that ICI treatment selectively impacts hippocampal-dependent cognitive function.

### Combined CTLA-4 and PD-1 blockade reduces synaptic density and myelination; and exaggerates CNS autoimmunity

To understand how combination ICI impairs cognitive function and further determine the extent of neuronal damage, we then investigated markers that are indicators of neuronal health, including pre- and post-synaptic density proteins (synaptophysin and PSD-95 respectively, **Fig. 3A-H**) and myelination using myelin basic protein (MBP, **Fig. 3I-J**). Both the Melanoma + Veh and Melanoma + ICI-treated mice showed a significant decline in the hippocampal synaptophysin immunoreactivity and therefore pre-synaptic density in the DG molecular (ML, **Fig. 3A, C**) and the CA1 stratum radiatum (SR. **Fig. 3B, D**) layers. For the post-synaptic density protein, PSD-95, we found a significant decrease in the DG ML **(Fig. 3E, G)** and CA1 SR **(Fig. 3F, H)** layers in the tumor-burdened groups with or without ICI treatment. These data indicate a significant synaptic loss post-ICI treatment. Cancer therapies, including cranial irradiation, have been shown to impact myelination in the brain [24] negatively. Therefore, we assessed myelination in the hippocampal corpus callosum using volumetric quantification of MBP immunoreactivity (**Fig. 3I-J**). All three treated groups, ICI, Melanoma + Veh, and Melanoma + ICI had significantly lower myelination than control-vehicle animals.

**Figure 3.**
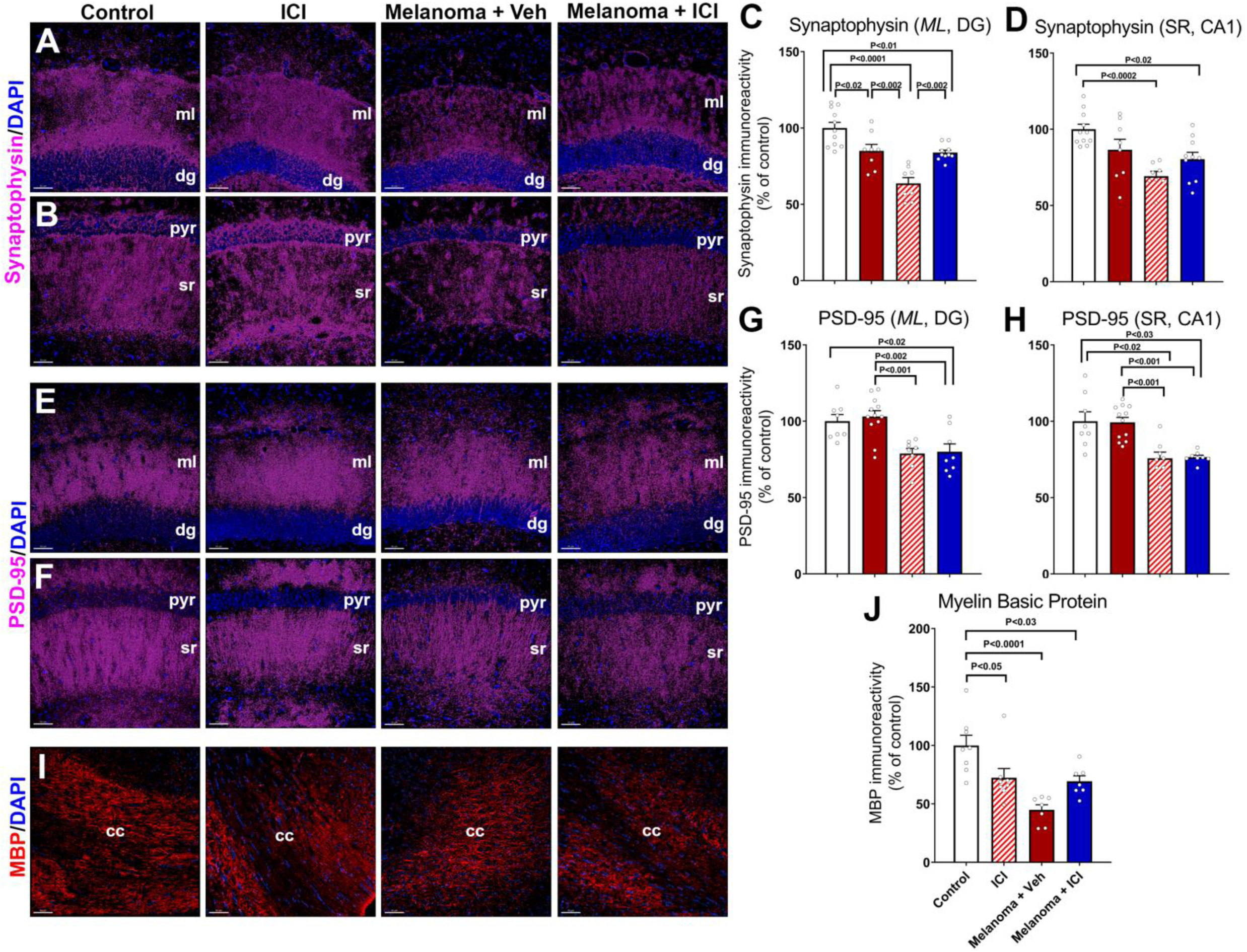
ICI treatment reduces synaptic integrity and myelination. Adult WT mice with or without melanoma received ICI (anti-CTLA4 and anti-PD-1) or isotype control (ITC) treatment for three weeks (as in Fig 1A). At eight weeks post-ICI treatment, mice were euthanized, fixed brains were collected for coronal cryosectioning, and immunostained for the pre- and post-synaptic markers and myelin basic protein (MBP). **(A-D)** ICI-treated control and cancer-bearing mice, and melanoma-bearing mice treated with vehicle showed significant reductions in the pre-synaptic protein, synaptophysin immunoreactivity (magenta, and DAPI nuclear stain) volume compared to controls in the molecular layer (ml) of the hippocampal dentate gyrus (*dg*, **A, C**), and stratum radiatum (*sr*) layer emanating from the CA1 pyramidal (pyr) neurons **(B, D)**. Data are presented as mean ± SEM (*N*=8 mice per group). *P* values were derived from ANOVA and Tukey’s post hoc test. Scale bars, 100 µm **(A)**, 50 µm **(B)**, and 5 µm **(d1)**.

To determine if ICI therapy modulates the CNS and creates a conducive state for autoimmune neuroinflammatory diseases, concurrently, we studied a mouse model of multiple sclerosis (MS) like disease, experimental autoimmune encephalomyelitis (EAE). The ICI group was treated with combination ICI as previously described above (anti-PD1, thrice weekly; anti-CTLA-4, twice weekly; for three weeks). One week after ICI therapy, control, and ICI mice were divided into two groups: active EAE and adoptive transfer (passive) EAE (**Suppl. Fig. S4 A**). Both formats of EAE revealed exaggerated clinical scores and severity in mice that received ICI, compared to control groups (**Suppl. Fig. S4 B-C**), thus supporting the hypothesis that ICI predisposes CNS to exaggerated autoimmune neuroinflammation. These results suggest that ICI is detrimental to two critical components of neuronal health: synaptic proteins and myelination.

### Immune checkpoint inhibition impairs neuronal plasticity and spares neurogenesis

To evaluate the neurodegenerative consequences of ICI treatment in cancer-bearing mice, we quantified neurogenesis using BrdU-NeuN dual-immunofluorescence by estimating the percentage of mature neurons differentiated from BrdU-labeled neural stem/progenitor cells. Briefly, after the completion of three weeks of ICI treatment, mice were treated with BrdU (50 mg/kg, once daily for 6 days) and euthanized about 8-weeks later. Frequencies of BrdU^+^ progenitor cells differentiating into mature neurons were comparable across all experimental groups **(Fig. 4A-B)** suggesting that neurogenesis is unaltered. To further determine the neuronal activity, plasticity-related immediate early gene (IEG) product cFos was co-stained with the mature neuronal marker (NeuN) in the DG blade. We found a significant decline in cFos-NeuN dual-labeled cells in the control brains treated with ICI and Melanoma + Veh mice brains (**Fig. 4C-D**, *P*<0.05 and *P*<0.003, respectively) indicating decline in the mature neuronal plasticity. Melanoma-bearing mice brains treated with ICI did not differ from controls.

**Figure 4.**
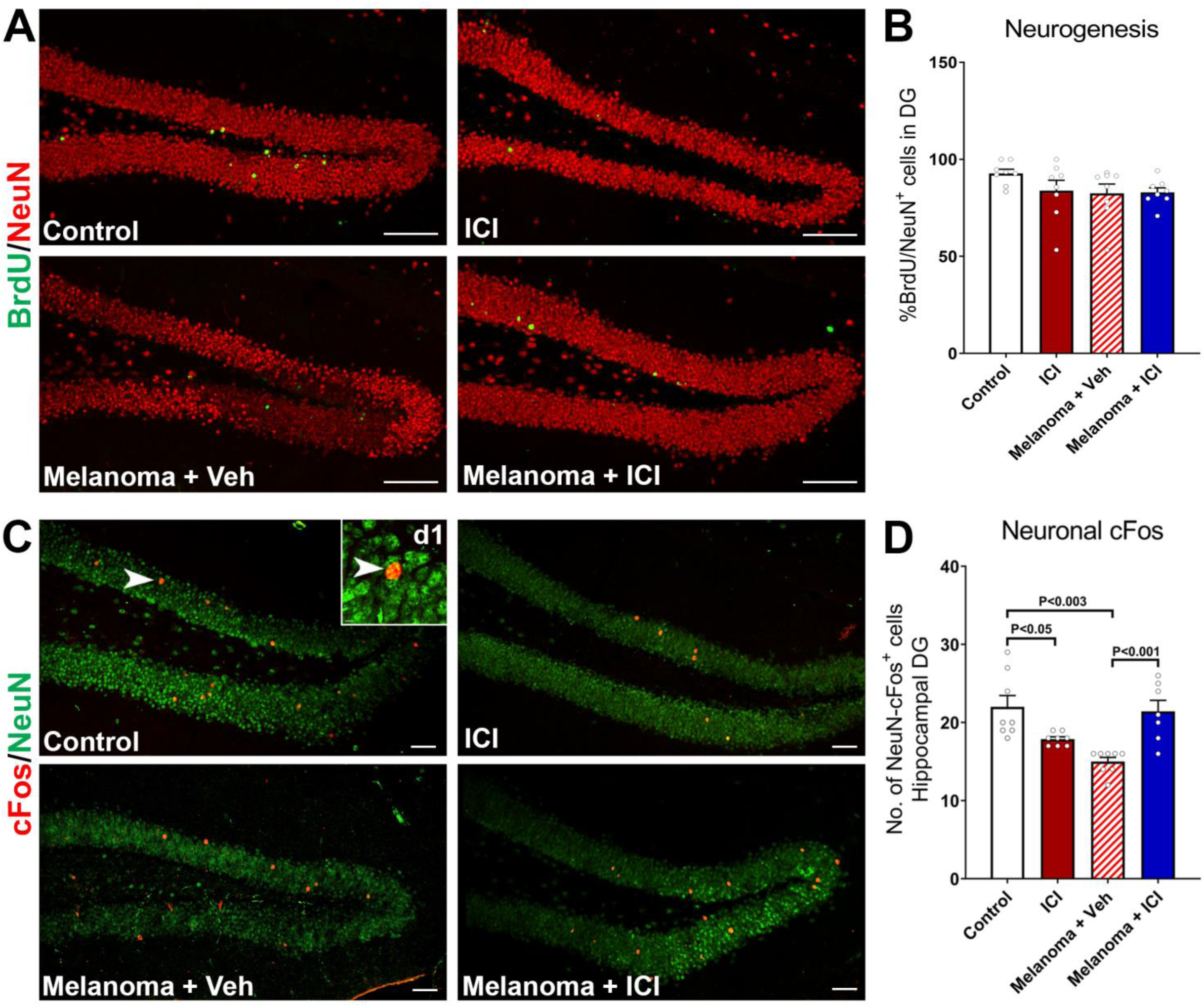
ICI treatment did not impact neurogenesis but reduced mature neuronal plasticity. ICI-treated mice with or without melanoma received a thymidine analog, BrdU injections 3 weeks before euthanasia to track the formation of new neurons from neural progenitor cells in the hippocampal sub-granular zone. Brains were collected 8 weeks post-ICI treatment and dual- immunostaining was conducted to determine the status of neurogenesis (BrdU-NeuN^+^) and neuronal plasticity (cFos-NeuN^+^). **(A-B)** The control and cancer-bearing mice with or without ICI treatment did not show significant differences in the percentage of BrdU^+^ neural progenitor cells differentiating into mature neurons (NeuN^+^) in the hippocampal dentate gyrus (DG). **(C-D)** Dual-immunofluorescence staining and 3D algorithm-based quantification of a neuronal plasticity-related immediate early gene product, cFos (red) and the mature neuron marker (NeuN, green) showed a significant decline in the number of cFos-NeuN^+^ dual-labeled cells in the control mice brains receiving ICI and melanoma-bearing mice brains receiving vehicle treatments. **(d1)** A high-resolution confocal z stack showing dual-labeled mature neuron (NeuN) expressing cFos. Data are presented as mean ± SEM (*N*=8 mice per group). *P* values were derived from ANOVA and Tukey’s post hoc test. Scale bars, 50 µm.

### CTLA-4 and PD-1 blockade elevate microglial activation but not astrogliosis

To determine the long-term effects of ICI on microglia and astrocytes, we measured the activation and hypertrophy status, respectively, using dual-immunofluorescence staining and laser scanning confocal microscopy of hippocampal slices at 8-weeks post-ICI treatment. The quantification of CD68, IBA1 immunoreactivity, and co-localization was facilitated by *in silico* 3D algorithm-based volumetric quantification of single and co-localized surfaces as described (**Fig. 5A**) (Markarian et al., 2021). The ICI-treated, melanoma-burdened mice treated with vehicle (Melanoma + Veh), as well as the melanoma-bearing mice receiving ICI (Melanoma + ICI) showed significant increases (*P*’s<0.0001) in CD68-IBA1 dual immunoreactivity in the hippocampal dentate gyrus (DG) indicating microglial activation **(Fig. 5B-C)**. These results align with earlier flow data showing higher levels of CD45^+^/CD11b^+^/MHCII^+^ cell population at 72 hrs (**Fig. 1I, K, N)**. Because, microglial activation is often accompanied by astrogliosis during cancer therapies (Acharya et al., 2016b; Markarian et al., 2021), we tested the effect of ICI on glial interplay by determining the volume of GFAP immunoreactivity (astrocytes) in the DG. We did not find significant differences in GFAP immunoreactivity in any of the ICI-treated or tumor-burdened groups compared to the control-vehicle group **(Fig. 5D-E)**. However, we observed a ∼20% decline (*P*<0.02) in GFAP immunoreactivity in the Melanoma + ICI group compared to the Melanoma + Veh (ITC)-treated group. These data suggest that ICI primarily triggers microglial activation while astrocytic hypertrophy is unaltered.

**Figure 5.**
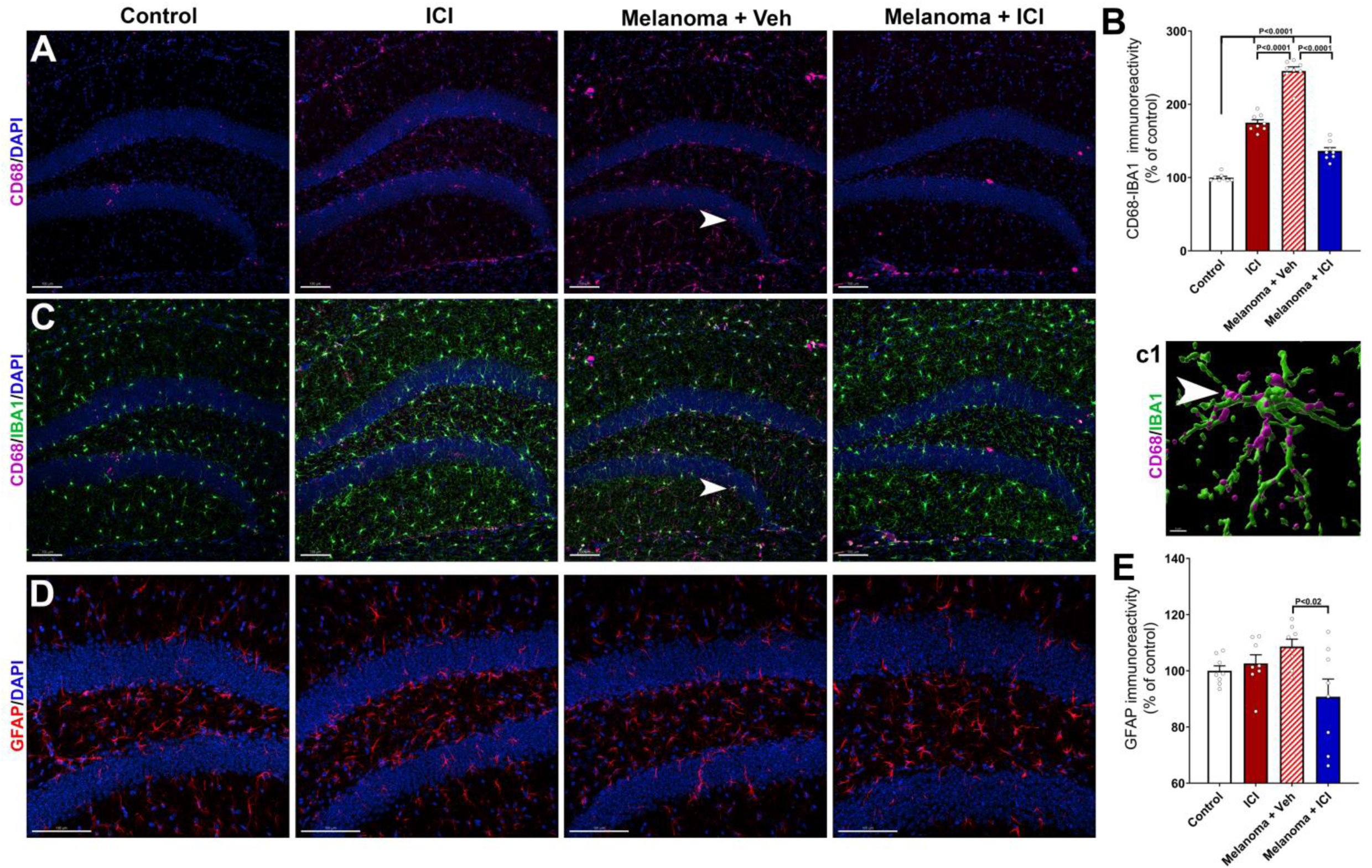
ICI treatment elevates microglial activation but not astrogliosis. Adult WT mice with or without melanoma received ICI (anti-CTLA4 and anti-PD-1) or isotype control (ITC) treatment for three weeks as in Fig 1A. At eight weeks post-ICI treatment, mice were euthanized, fixed brains were collected for coronal cryosectioning, and immunostained for the activated microglia and astrocyte markers. **(A-C)** 3D algorithm-based volumetric quantification of lysosomal marker (CD68, magenta, **A**) and its colocalization with the pan microglial maker (IBA1, green) in the hippocampal dentate gyrus and dentate hilus (DAPI, blue, nuclear stain, **A, C**) show significant elevations in the CD68-IBA1 co-labeling in the control and melanoma-bearing mice brains treated with ICI, and melanoma-bearing mice receiving ITC treatment. **(c1)** A representative high-resolution immunoreactive surface rendering of CD68^+^ puncta (magenta) with IBA1+ microglial cell surface is shown. **(D-E)** ICI treatment in the control or melanoma-bearing mice did not elevate astrocytic immunoreactivity (GFAP, red, DAPI, blue, nuclear stain) in the control and melanoma-bearing mice brains. Melanoma + ICI group showed a decline in the GFAP immunoreactivity compared to the Melanoma + Vehicle group. Data are presented as mean ± SEM (*N*=8 mice per group). *P* values were derived from ANOVA and Tukey’s post hoc test. Scale bars, 100 µm **(A, C, D)**, and 5 µm **(c1)**. **(E-H)** Concurrently, 3D algorithm-based volumetric analysis of post-synaptic density, PSD-95 immunoreactivity (magenta, and DAPI nuclear stain) showed significant reductions in the hippocampal *ml* **(E, G)** and *sr* **(F, H)** subregions in melanoma-bearing mice brains ± ICI treatment and control mice treated with ICI compared to the vehicle-treated controls. **(I-J)** Volumetric quantification of MBP immunoreactivity (red, and DAPI nuclear stain) in the white matter (corpus callosum, *cc*, **I**) showed significant reductions in all the groups including control treated with ICI, melanoma and melanoma mice received ICI treatment compared to controls (**J**). Data are presented as mean ± SEM (*N*=8 mice per group). *P* values were derived from ANOVA and Tukey’s post hoc test. Scale bars, 50 µm **(A-B, E-F, I).**

## DISCUSSION

The core finding of our study supports our hypothesis that combinatorial blockade of CTLA-4 and PD-1 to treat melanoma elicits immune-related adverse effects on the brain physiology by inducing neuroinflammation (microglial activation) that culminates in neurodegeneration (synaptic loss), ultimately resulting in a decline of cognitive function.

Emerging clinical observations are increasingly reporting neurocognitive complications accompanied by neurotoxicities following ICI therapy (Duong et al., 2021; Fleming et al., 2023; Myers et al., 2023) in patients with melanoma, lung cancers, and Merkel-cell carcinoma at five weeks post-therapy (Duong et al., 2021). Rogiers and colleagues reported long-term neurocognitive complications in about 41% of metastatic melanoma survivors up to 2 years after ICI treatment (Rogiers et al., 2020a; Rogiers et al., 2020b; Rogiers et al., 2023). These clinical studies suggest that ICI treatment-induced neurotoxicity and neurocognitive decline are progressive, and clinical symptoms appear within weeks post-ICI and last several years post-treatment. Our past clinical and preclinical studies and others have shown a relationship between proinflammatory cytokines and CRCI following chemo- and radiation therapy (Allen et al., 2020; Cheung et al., 2015; Markarian et al., 2021; Oppegaard et al., 2021). Chronic cytotoxic chemotherapy (cyclophosphamide, and doxorubicin) significantly impair performance on hippocampal and cortex-dependent cognitive tasks (Acharya et al., 2015; Allen et al., 2019; Christie et al., 2012; Usmani et al., 2023), and these deficits were associated with reduced neurogenesis, elevated neuroinflammation, and a significant decline in synaptic density. Elevated neuroinflammation underlies many, if not all, neurodegenerative disorders in humans, and our data suggest that neuroinflammation is one of the major contributory factors for long-term CRCI. The upregulation in proinflammatory cytokine signaling can mediate neuroinflammation thereby leading to cognitive dysfunction. Elevated neuroinflammation in CRCI is a significant avenue to pursue with the advancement of novel cancer therapeutics including ICI.

### Neuro-immune homeostasis is altered during combination ICI treatment

ICI treatment could exaggerate neurocognitive impairment by increasing peripheral immune response leaking into CNS or triggering neuroinflammation within the brain (McGinnis and Raber, 2017). Our comprehensive assessment of immune cells in the CNS using flow cytometry provides a window into the cellular mechanisms of IRAE for combination ICI. Elevated numbers of lymphoid cells (CD4 T, CD8 T, and B cells) with a memory phenotype (CD27^+^) indicate autoreactive lymphocytes unleashed in the brain post-ICI treatment. While it remains to be determined whether ICI activates CNS resident memory cells or promotes infiltration from the periphery, elevated lymphocytes could elicit their effector functions in the brain and drive a neuroinflammation-like microenvironment. Until recently, CNS was considered an immune-privileged site. However, it is increasingly becoming evident that CNS and the peripheral immune system are constantly communicating (Kipnis, 2016; Salvador and Kipnis, 2022). Others have recently shown that ICI antibodies can enter the brain (Vinnakota et al., 2024), which could release the breaks on the CNS tissue resident lymphoid cells. A modestly higher number of monocytes and comparable NK cells suggest local alteration of the immune cell rather than peripheral infiltration, which is the primary driver of observed changes in the immune cells in the brain. Our plasma cytokine data suggest significantly elevated pro-inflammatory cytokines including IL1α, TNFα, IL6, IL10, IL17, and IL12. Previously, using a rodent model of radiation-induced cognitive dysfunction, we have shown elevated pro-inflammatory cytokines that were accompanied by microglial activation and reduced synaptic density (Markarian et al., 2021). IL1α and TNFα, in association with CNS complement cascade factors (C1q and C3), were found to be neurotoxic leading to synaptic loss (Liddelow et al., 2017). Elevated IL6, IL10, IL17, and IL12 post-ICI treatment are also indicators of elevated inflammation. ICI-mediated neuroinflammation is further supported by an overall decline in the percentage and number of CD11b^+^ cells in the ICI-alone group and marginally higher frequencies of MHCII-positive myeloid cells. Thus, ICI treatment-induced elevation of lymphocytes may trigger subsequent inflammatory reactions in the brain, leading to CNS neuroinflammation and neuronal damage.

### Neuroinflammation post-ICI treatment

Combinatorial ICI therapy using anti-CTLA4 and anti-PD1 controlled melanoma growth *in vivo* (**Fig. 1B**). This model allowed us to study the impact of such targeted cancer therapy on brain function. Our past studies, using chemotherapy and radiation therapy rodent models (Acharya et al., 2016a; Acharya et al., 2011; Acharya et al., 2016c; Acharya et al., 2015; Allen et al., 2019; Markarian et al., 2021; Montay-Gruel et al., 2019), have shown elevated neuroinflammation, particularly microglial activation, in the brain, leading to cognitive impairments. We found that combination ICI led to a significant increase in microglial activation **(Fig. 2)** in mice with or without cancer, as shown by elevated hippocampal dual- immunoreactivity for CD68 and IBA1 at 6-8 weeks post-ICI treatments. We did not find a significant increase in astrogliosis *in vivo* **(Fig. 2)**. IBA1 is a pan microglial marker expressed by both resting and activated microglia, whereas CD68 is a lysosomal protein marker expressed by phagocytotic microglia and macrophages. Long-term elevation in the CD68-IBA1 dual- immunoreactivity following cancer induction and ICI treatment indicates proinflammatory (phagocytotic) activation of microglia that could lead to a detrimental impact on neuronal and synaptic health. Microglia play important roles in the clearance of dead cells and sub-cellular debris and synaptic pruning *in vivo* (Paolicelli et al., 2011; Paolicelli et al., 2022). Proinflammatory activation of microglia may lead to excessive synaptic pruning and neuronal damage, leading to compromised neuronal and neurocognitive function.

### Neuron plasticity and myelin loss post-ICI treatment

We evaluated surrogate markers of neuronal health including myelin (MBP), pre- and post-synaptic density proteins (synaptophysin, and PSD-95 respectively) and neuronal plasticity IEG marker, cFos-NeuN. No-cancer mice receiving ICI, cancer-bearing mice with or without ICI treatment showed a significant decline in the myelin basic protein immunoreactivity **(Fig. 4)**. Both, infiltration of immune cells and resident immune cell (microglia) activation may lead to myelin damage. Our results from the active model of EAE suggest that ICI pretreatment exposes CNS to exaggerated CNS autoimmunity (**Suppl. Fig. S4**). To rule out the effect of ICI on the peripheral immune system during active EAE, we used an adoptive transfer EAE model. Accordingly, we also observed similar results of exaggerated EAE, strengthening our hypothesis that an ICI-mediated neuroinflammation-like state is conducive for autoreactive lymphocytes to orchestrate higher levels of damage to myelin and ultimately drive neural degeneration. Previously, using an autoimmune model (EAE), we have shown that infiltration of activated, inflammatory immune cells including T cells, NK cells, and B lymphocytes trigger a huge myelin loss and axonal degeneration *in vivo* (Othy et al., 2020). We found elevated expression of MHC class II positive cells and CD68-IBA1 positive activated microglia in the brain following ICI treatment (**Fig. 1**, and **5**). Chronic elevated neuroinflammation has been linked with long-term myelin loss and progressive neurodegeneration *in vivo* (Loane et al., 2014). Indeed, we found a loss of synaptic integrity in the hippocampal molecular layer (ML) and CA1 stratum radiatum of the dentate gyrus (DG), important for the learning and memory processes. ICI treatment to intact or melanoma-bearing mice and melanoma burden itself led to a significant decline in the pre-synaptic protein, synaptophysin, at two months post-ICI treatment (**Fig. 3**). Synaptic loss was more pronounced in melanoma-bearing mice receiving vehicle or ICI treatments for the post-synaptic density protein PSD-95 (**Fig. 4**). Synaptophysin plays an essential role in per-synaptic vesicular trafficking, and its loss is often associated with neurodegenerative conditions, including Alzheimer’s disease (Evans and Cousin, 2005; Glantz et al., 2007). We have also shown reduced synaptophysin levels in the irradiated brains coincident with cognitive decline (Markarian et al., 2021; Parihar et al., 2014). Post-synaptic density protein, PSD-95, is a scaffolding protein that plays essential roles in excitatory neurotransmitter receptor clustering and function (Glantz et al., 2007; Keith and El-Husseini, 2008). ICI treatment-induced myelin and synaptic protein loss could inadvertently impact neuronal health and neurotransmission. We found reduced expression of an immediate early gene (IEG), cFos, within the mature neurons (NeuN^+^) following ICI treatment. Neuronal cFos expression is linked with neuronal activity, particularly voltage-gated calcium channel-mediated calcium influx via neuronal NMDA receptors (Kang et al., 2001). cFos facilitates hippocampal neuronal LTP and long-term memory formation. We found reduced neuronal cFos (cFos-NeuN^+^) in the intact mice receiving ICI treatment and melanoma-bearing mice brains (**Fig. 3**). Conversely, hippocampal neurogenesis (BrdU-NeuN), the formation of new neurons from neural stem/progenitor cells, was unchanged (**Fig. 4**). Collectively, ICI- and proinflammatory environment-induced myelin, pre- and post-synaptic protein loss, and reduced neuronal plasticity *in vivo* indicate a long-term detrimental impact on neuronal function that could culminate into neurocognitive impairments.

### ICI-induced cognitive dysfunction

Melanoma-bearing and intact mice receiving ICI treatment underwent behavior and cognitive function testing two months post-treatment. The open field test (OFT) and anxiety-related light-dark box (LDB) test did not show increased anxiety in cancer-bearing mice (**Fig. 5B-C**), indicating an absence of neophobic behavior during cognitive function testing. ICI-treated no-cancer mice showed reduced time spent in the open zone (OFT) and reduced latency in entering the dark compartment. Thus, ICI-treated mice were less explorative due to a marginal increase in anxiety. This resulted in them spending more time in the peripheral zones of the open arena (OFT) or the dark compartment (LDB). Subsequently, mice were tested on a hippocampal-dependent object location memory (OLM) task that determined spatial context memory (**Fig. 5D-E**). ICI treatment in either cancer or no-cancer mice led to a significant decline in the performance of the OLM task. Lastly, mice were trained on fear conditioning, fear extinction and tested on the hippocampal- and medial prefrontal cortex (mPFC)- dependent fear extinction memory tests (**Fig.5F-H**). During the conditioning phase (three tone-shock pairs), all groups of mice showed progressively increased freezing. No-cancer mice receiving ICI treatment showed the highest freezing compared to controls (P<0.01). This behavior is related to their increased anxiety levels, as observed in the OFT and LDB tasks. For the next three days, all groups of mice received fear extinction training (repeated tones without shocks) in the same spatial context and the odor cue 24 hours apart (**Fig. 5F**). All groups of mice showed reduced freezing, except for no-cancer ICI-treated mice that remained frozen and did not learn to extinct the aversive memory. At 24 hours post-extinction training, ICI treatment to cancer or no-cancer mice during the extinction test led to a significantly elevated freezing, indicating a compromised hippocampal-dependent fear memory-consolidation process (**Fig. 5G**). Melanoma-bearing mice receiving vehicle did not differ from controls. At 24 hours after the extinction test, the context and the odor cue within the fear testing arena were changed to engage mPFC. In this test, no-cancer mice receiving ICI remain frozen compared to controls **(Fig. 5H)**. Thus, ICI treatment in melanoma-bearing mice led to a long-term disruption of hippocampal-dependent learning and memory and memory consolidation processes *in vivo*.

### Limiations of the study

Our study provides pre-clinical evidence that combined treatment with anti-CTLA-4 and anti-PD-1 is neurotoxic, leading to an elevated neuroinflammatory environment and thus causing neurocognitive decline. We showed this using pre-clinical cancer-free and melanoma-bearing mouse models. One of the studies showed that anti-PD-1 can enter the CNS and interact with microglia leading to pro-inflammatory activation (Vinnakota et al., 2024). This possibility is yet to be determined in our model using combined ICI treatment. We also observed significant elevations in T cells, including memory T cells, in the ICI-treated brains. Determining the CNS immune modulatory role of memory T cells in the context of microglial activation could shed light on sustained neuroinflammation in the brain exposed to ICI. A long-term assessment of cognitive function beyond one month post-ICI treatment would allow studying long-lasting neurotoxic effects of ICI that may provide a window of opportunities to develop neuroprotective strategies.

***Conclusion:*** Immunotherapy for cancer elevates peripheral immune cell infiltration and activation in the brain that triggers neuroinflammation, myelin and synaptic loss, reduces neuronal plasticity, and significantly impacts hippocampal-dependent cognitive function. Our preclinical observations using a melanoma-bearing mouse model treated with ICI closely model human cancer survivors. Emerging human studies report detrimental neurocognitive sequelae of ICI in at least 41% of cancer survivors without available clinical recourse (Duong et al., 2021; McGinnis and Raber, 2017; Rogiers et al., 2020a; Rogiers et al., 2020b). Our study points to elevated inflammation within the brain that promotes neurodegenerative consequences. We observed similar long-term neuroinflammation in chemotherapy- and cranial radiation therapy-exposed brains, leading to cognitive impairments (Acharya et al., 2016a; Acharya et al., 2011; Acharya et al., 2016c; Acharya et al., 2015; Allen et al., 2019; Markarian et al., 2021; Montay-Gruel et al., 2019). Based on the foregoing, we propose that ICI-induced peripheral immune cell infiltration and neuroinflammation snowball into progressive neuronal and synaptic damage, leading to cognitive dysfunction. Thus, mitigation strategies targeted to thwart or reduce neuroinflammation without compromising anti-cancer efficacy will provide a therapeutic benefit to prevent cognitive decline and improve the quality of life for thousands of cancer survivors receiving Immunotherapy.

## Supporting information

All_Supplemental_Figures

## Acknowledgments

This research was supported by the National Institutes of Health (NIH) awards (R01CA251110, R01CA262213) to M.M.A, R01CA251110-03S1 to O.V.I. (PI, M.M.A.), UC Irvine Chao Family Comprehensive Cancer Center (CFCCC) Pilot Award (M.M.A.), HESI-Thrive award (Health and Environmental Sciences Institute), and American Brain Tumor Association (ABTA) Discovery (DG2000029) awards to M.M.A, R01AI168063 to S.O (PI), and U01AI160397 to S.O (Co-I). We thank Dr. F. Marangoni (UC Irvine) for providing assistance in developing the melanoma model. We thank Dr. A. Agrawal, Dr. J. E. Baulch, Robert P. Krattli, Jr., and Tracy Nguyen for their technical assistance. We also thank the support of the UCI CFCCC Genomics Research and Technology Hub shared resource supported by the National Cancer Institute (NIH, award P30CA062203). The content is solely the responsibility of the authors and does not necessarily represent the official views of the National Institutes of Health.

## Author Contributions

Conception and design: SO & MMA. Study supervision: SO & MMA

Development of methodology: OVI, AD, SMK, & AZ Acquisition of data: OVI, AD, SMK, NHH, & AZ

Analysis and interpretation of data: OVI, AD, SMK, SO, & MMA Administrative, technical, or material support: OVI, AD, SMK, & MMA Writing, review and/or revision of the manuscript: OVI, AD, SO, & MMA

## MATERIALS AND METHODS

Detailed materials, methods and protocols are provided in the **Supplemental Methods and Materials** section.

### Study Design

This study aimed to determine the effects of immune checkpoint inhibitors (ICI) on cognitive function of melanoma mouse models. Wild-type, immunocompetent, male mice (aged 10-12 weeks) were randomly divided into 4 groups (N=12/group): Control mice did and did not receive ICI (Con ± ICI); melanoma mice did and did not receive ICI (Mel ± ICI). Melanomas were induced via intradermal subcutaneous using the D4M-3A.UV2 cells line and then received combination ICI treatments (anti-PD1 and anti-CTLA-4; BioXcell) one week post-cancer induction. The ICI treatments were administered for 3 weeks via intraperitoneal injections. Mice that did not receive ICI treatments were given isotype-matched antibodies. Neurogenesis in the brain was accessed by injecting BrdU (5-Bromo-2′deoxyuridine, Sigma) for 6 days post ICI-treatments. One month following the initial ICI treatment, cognitive behavior tests were performed, followed by transcardial perfusion and collection of brain tissues for downstream analyses (immunohistochemistry and flow cytometry).

### Animals, tumor induction, and tissue analysis

All animal use procedures and safety protocols were approved by UCI Intuitional Animal Care and Use Committee and were consistent with NIH guidelines. Adult (8-10 week) male WT mice were randomly divided into four groups: non-tumor burden mice treated with vehicle, or combinatorial (CTLA-4 and PD-1) ICI and melanoma (D4MUV2) burden mice treated with vehicle or combinatorial (CTLA-4 and PD-1) ICI. Melanoma tumor induction was accomplished via bilateral, subcutaneous abdominal tent injection of 5 × 10^5^ syngeneic D4MUV2 cells. Tumors were allowed to engraft for one week before starting the ICI treatment. Anti-CTLA-4 and anti-PD1 were diluted with the dilution buffer (InVivoPure, pH 6.5, (BioXCell) to the dosage of 1mg/mouse anti-CTLA-4, and 200µg/mouse anti-PD1. Anti-CTLA-4 were given intraperitoneal injections 3 times per week (every other day), and anti-PD1 for 2 times/week every third day. ICI treatments were given for 3 weeks. Control mice that did not receive the combined ICI treatments were injected with isotype-matched control (ITC) antibodies (Armenian hamster IgG, and Rat IgG2a, BioXCell) following the same injection schedule as the mice in the treatment groups. For the flow cytometry study, at 72 hours after the last ICI injection, animals were euthanized and brains were collected for the analysis. For the cognitive function study, mice after completion of ICI treatment, mice were allowed to recuperate for one month before undergoing cognitive function tasks for 3-4 weeks. Mice were also injected with BrdU (50 mg/kg, IP, once daily for 6 days) to study neurogenesis 72 hours after completion of ICI treatments. Mice were sacrificed 48 hours following the completion of behavior. Brains were collected after transcardial perfusion, fixed using 4% paraformaldehyde and cryo-sectioned (30 um, coronal). For gene expression analysis, the micro-dissected hippocampus was flash frozen at -80C.

### Behavior testing

Mice were randomly divided into control-vehicle, melanoma-ITC (vehicle), control-ICI, and melanoma-ICI groups. Our past studies have shown that the group size of 8-10 mice/group for behavior and 4-6 mice/group for the molecular or tissue analyses achieved sufficient power (α=0.05) to reach the statistical significance (references). 1-month post ICI treatment, mice underwent light-dark box and open field testing to assess anxiety, novel place recognition (NPR) to assess spatial memory. For the light-dark box (LDB) test, the time spent in light and dark chamber and number of transitions between compartments was assessed. The mice then underwent open field test (OFT) which evaluates an animal’s partiality for staying in the open center of the field (30% of arena) or the shaded outer edge (70% of the arena). The NPR task depends on animal’s inherent preference to explore a novel location of an object and relie on the intact hippocampal function (32,33). Time spent interacting with the familiar versus novel object or location was recorded and the data calculated as the Memory Index (MI: [Novel location or Novel object exploration time/Total exploration time] – [Familiar location exploration time/Total exploration time]) × 100. The same cohort of mice also underwent a fear extinction memory test as described (see Supplemental Information). All tests were scored by blinded observers to avoid experimental group bias. After conclusion of behavior testing, mice were euthanized via transcardial perfusion and brains were fixed using 4% PFA for the immunohistochemistry (IHC) analyses.

### Immune cell isolation from the brain and flow cytometry

Please see the Supplemental Information for details about the brain cell isolation protocol. This FACS analysis was carried out using a Novocyte Quanteon (Agilent Technologies, California, US).

### Dual-immunofluorescence staining, confocal microscopy, and 3D algorithm-based volumetric quantification

The PFA-fixed brains from each treatment group underwent dual-immunofluorescence staining (2 sections/brain and 4 brains/group) including GFAP, IBA1, synaptophysin, PSD95, myelin basic protein (MBP), BrdU-NeuN, NeuN-cFoS, and IBA1-CD68. Imaris modules were used to blindly and unbiasedly 3D deconvolute and volumetrically quantify neuronal and glial surfaces. For more details regarding the primary and secondary antibodies, immunostaining, and *in silico* modeling, please see the Supplemental Information section.

## Data analysis

Statistical analyses were performed to confirm overall significance (GraphPad Prism, v8.0). For the analysis of ICI treatment to the no-cancer and cancer-bearing mice, two-way ANOVA or repeated measures ANOVA, and Bonferroni’s multiple comparisons test were performed. For the tumor studies, P values were derived from the Mann-Whitney *U* test or long-rank test for survival studies. All results are expressed as the mean values ± SEM. All analyses considered a value of P≤0.05 to be statistically significant.

## SUPPLEMENTAL MATERIALS AND METHODS

### Lead Contact

Further information and requests for resources and reagents should be directed to and will be fulfilled by the lead contact, Munjal Acharya.

### Materials Availability

This study did not generate any new unique reagents.

### Data and Code Availability

● All data reported in this paper will be shared by the lead contact upon request.
● This paper does not report original code.
● Any additional information required to reanalyze the data reported in this paper is available from the lead contact upon request.

## EXPERIMENTAL MODEL AND ANIMALS DETAILS

### Mice

All animals used in this study were in accordance with the National Institute of Health (NIH) guidelines and approved by the Institutional Animal Care and Use Committee (IACUC) at University of California, Irvine. Wild-type male mice (C57BL/6J, Jackson Laboratory RRID:IMSR_JAX:000664), aged 10-12 weeks were group housed (2-4 mice per cage) in standard conditions (20 °C±1 °C; 70% ± 10% humidity;12h:12h light and dark cycle) and given a standard rodent chow diet (Envigo Teklad 2020X) by the University Laboratory Animal Resources (ULAR).

### D4M-3A.UV2 Melanoma induction and tumor measurement

D4M-3A.UV2 mouse melanoma cells were cultured in monolayer with Gibco^TM^ DMEM (1X) + GlutaMAX^TM^-I (FisherSci, Cat. 10-569-010) supplemented with 10% Gibco^TM^ FBS (FisherSci, Cat. 10-082-147) under 37°C/5% CO2 conditions. On the day of tumor induction, D4M-3A.UV2 cells were trypsinized in Gibco™ TrypLE™ Express Enzyme (FisherSci, Cat. 12-605-010) and centrifuged (300g, 5 minutes). Trypsin was neutralized with DMEM + 10% FBS, washed, and centrifuged (300g, 5 minutes) to remove excess TrypLE. Cells were resuspended in Gibco™ Hibernate™-A Medium (FisherSci, Cat. A1247501) and kept on ice until tumor induction. 1X PBS (FisherSci, Cat. 14-190-144) was used to spray down the injection area (left flank) to help the researcher detect the injection site without interference from the fur. Each mouse subject received an intradermal injection dose of 1 x 10^6^ D4M-3A.UV2 cells in 0.2mL Hibernate Medium (FisherSci) on the left flank using a 31 Gauge insulin syringe (BD Sciences, Cat. 328411-1) until a small bleb appeared underneath the skin. The bleb indicates that the cells were successfully injected under the dermis. Mice were monitored every other day for tumor growth. Tumor growth was measured using an electronic caliper along the tumor’s horizontal and vertical axis. The maximum diameter was determined as the larger diameter between vertical and horizontal measurements. The tumor volume was calculated using the formula 0.5 x *a x b*^2,^ where *a* is the minimum diameter and *b* is the maximum diameter. Mice were treated with Immunotherapy when the tumor volumes were approximately 20-30 mm^3^ (7 days post-tumor induction).

### Combined Immunotherapy of anti-CTLA-4 and anti-PD1

After one week of melanoma induction, mice in the treatment groups received the combined Immunotherapy of anti-CTLA-4 (BioXcell, Cat. BP0032 RRID: AB_1107598) and anti-PD 1 (BioXCells, Cat. BP0273 RRID: AB_2687796). Anti-CTLA-4 and anti-PD1 were diluted with InVivoPure pH 6.5 Dilution Buffer (BioXCells, Cat. IP0065) to 1 mg/mouse and 200µg/mouse, respectively. Anti-CTLA-4 was given intraperitoneal injections 3 times/week on Monday, Wednesday, and Friday, and anti-PD1 2 times/week on Monday and Thursday continuously for 3 weeks. Control mice that did not receive the combined ICI treatments were injected with isotype-matched control (Isotype control for CTLA-4: Armenian Hamster IgG, BioXcell, Cat. BP0091, RRID: AB_1107773; Isotype control for PD1: Rat IgG2a, BioXCell, Cat. BP0089, RRID: AB_1107769) following the same injection schedule as the mice in the treatment groups.

### Cell Isolation from Brain Tissue for Flow Cytometry

72 hours following the last ICI injection, mice brains were collected. Mice were placed in a CO2 chamber until respiration stopped and immediately perfused (intracardial) with 25ml of 1X PBS (Gibco, Cat# 10010-023) containing 2mM EDTA (Invitrogen, Cat# 15575-038) using 50ml syringes (BD Biosciences, Cat# 309653) and 21 ¾ gauge butterfly needle (Abbott Lab, Cat#4492). Brains were dissected and placed in 1 ml of AIMV media (Thermo Fisher, Cat. 12055083) in 15 ml conical tubes on ice (Falcon, Cat# 352097). The brain tissue was homogenized until all large chunks were broken apart using a 15 ml homogenizer (VWR, Cat# 47732-446). Samples were then transferred to a 15 ml vented cap conical tube (Celltreat, Cat# 229471) for enzymatic digestion using 0.5 mg/ml Collagenase IV (Gibco Cat# 17104-019), and 25 µg/ml DNase I (Thermo Scientific, Cat# J62229.MC) in AIM V media (5 ml per brain). Samples were incubated at 37°C while shaking (500 RPM) for 50 minutes. FBS was added to stop enzymatic activity, and cells were passed through a 100uM filter (Miltenyi, Cat# 130-098-463) and washed once with RPMI (Gibco, Cat# 21870-076 500ml) + 10% FBS (Omega Scientific, Cat# FB- 12) and transferred to the new conical tube. Myelin was removed from the samples using a 37.5% Percoll gradient centrifugation as described previously (Othy et al., 2020). The samples were resuspended in 1ml of 37.5% Percoll and then filled to the 7.5ml mark with more of the 37.5% Percoll. Tubes were slowly overlayed with 1X PBS up to the 10ml mark using a transfer pipette (Fisherbrand, Cat# 13-711-7M) before centrifuging for 25 minutes at room temperature at 2300 RPM with the brakes off. Widened 1000ul pipette tips (Biotix, Cat# 63305410) were used to remove the myelin layer and supernatant. The pellet was resuspended in 1ml of RPMI+ 10% FBS. and transferred to a new 15ml conical tube, filled up to 10ml with RPMI+ 10% FBS, and centrifuged for 5 minutes at 1800 RPM at 4°C. RBS lysis was performed using ACK lysis buffer 0.15M NH4CL (Sigma Aldrich, Cat# A-0171), 10MM KHCO3 (Fisher Scientific, Cat# P184-500), 0.1mM NaEDTA (Sigma Aldrich, Cat# E-1644). RBC lysis was stopped by restoring the osmolarity with PBS and 10% FBS. Cells were then stained for flow cytometry.

### Flow Cytometry

Samples were stained at 4°C, protected from light, and washed with FACS buffer between steps. Cells were first incubated with Fixable Viability Dye eFluor™ 780 (ThermoFisher, Cat#65-0865- 18) at 1:2000 dilution in PBS for 10 minutes. Fc receptors were blocked with TruStain FcX™ (anti- mouse CD16/32, Biolegend Cat# 101320) for 20 minutes then stained with following antibodies: VioletFluor™ 450 anti-mouse CD19 (Tonbo Biosciences, Cat# 75-0193-U100), BV480™ Rat anti-mouse I-A/I-E (MHCII) (BD Biosciences, Cat# 566086), BV 570™ anti-mouse Ly-6C (Biolegend, Cat# 128030), BV605™ anti-mouse/human CD11b (Biolegend, Cat# 101237), BV 650™ anti-mouse CD45.2(Biolegend, Cat# 109836), PE anti-mouse/human CD3e (Invitrogen, Cat# 12- 0031-83), FITC Anti-Mouse CD4 (Tonbo Biosciences, Cat# 35-0041-U100), PerCP Anti-Mouse CD8a (Tonbo Biosciences, Cat# 67-0081-U100), PE/Fire™ 700 anti-mouse NK-1.1 (Biolegend, Cat# 108774), and Alexa Fluor® 700 anti-mouse/rat/human CD27 (Biolegend, Cat# 124240). UltraComp eBeads™ Compensation Beads (ThermoFisher Cat# 01-2222-42), ArC™ Amine Reactive Compensation Bead Kit (ThermoFisher, Cat# A10346) and cells from spleen served as controls for compensation and single stained samples for gating. Flow data was acquired on a Novocyte Quanteon flow cytometer, which was analyzed using Novocyte’s NovoExpress software and FlowJo.

### Open Field Test and Object Location Memory

Four weeks after the initiation of the combined ICI treatment, behavior testings were administered to determine the impact of the combined ICI treatment on cognitive function. Open field test (OFT) and object location memory (OLM) test were performed. For both tasks, the experimental setup includes a strictly controlled room with appropriate lighting (50-70 lux), four square open-field arena boxes (30 x 30 x 30 cm), camera recording equipment (Noldus), and tracking software (EthoVision XT 17, Noldus). The OFT task evaluated an animal’s preference to stay in the open field (30% of the central zone) or the shaded outer edge (70% of the outer edge of the arena) and was performed on the first day of habituation of the OLM task. The time that the mice spent either in the open field versus the shaded area of the arenas was quantified to assess their exploratory behaviors. Spatial recognition and memory were evaluated using the OLM task, which highly depends on an intact and functioning hippocampus (2,3). The OLM tasks were conducted as reported previously (1, 7). Briefly, the mice were habituated with the testing room and open field arena boxes (30 x 30 x 30 cm) that had blue tape down the inside wall of the arena box. The boxes had a thin layer of bedding and no toys were present; the mice were allowed to familiarize themselves with the testing environment for 10 minutes per run in 3 consecutive days (with OFT data was collected on the first day of habituation). During test day, the familiarization phase was conducted by allowing the mice to explore two identical plastic toys that were magnetically secured in place 16 cm apart from each other for 5 minutes (familiar phase). After the 5-minute of familiarization, the mice were returned to the home cage for 5 minutes while the objects were cleansed with 10% ethanol and thoroughly dried. One object was then moved to a new location inside the box (novel place), 16 cm from the opposite corner of the other toy, which remained in its former spatial location (familiar place). Mice were returned to arena boxes and allowed to explore for 5 minutes. Following the test phase, all mice were returned to their home cage. Mice behavior from all phases was recorded using the Noldus camera system. The "head direction to zone" function from Ethovision software was utilized to track the mouse behavior and record their exploration time. Furthermore, to ensure unbiased analysis, time spent interacting (nose within 2 cm) with familiar versus novel place objects was scored by researchers blind to the experimental conditions. The discrimination index (DI) was calculated for each animal using the equation: ([Novel object or location exploration time/Total exploration time] – [Familiar object or location exploration time/Total exploration time]) × 100.

### Light-Dark Box (LDB)

After the OLM test day, mice anxiety-like behavior were evaluated by performing the light-dark box (LDB) test using established methods as reported previously (4, 8, and 9). The LDB arena consisted of a dark compartment (15 x 10 x 27 cm, 4 lux) connected to a light compartment (30 x 20 x 27 cm, 915 lux) via a small opening (7.5 x 7.5 cm). This task juxtaposes a mouse’s inclination to explore new environments with their level of anxiety to be in a well-lit space. Mice were allowed to explore in the LDB arena for 10 minutes, the amount of time spent in each compartment and the number of transitions between the light and dark compartments were recorded.

### Fear Extinction Memory

After OFT and OLM cognitive task, we performed fear extinction behavior reliant on hippocampal function to determine if the combined treatment of anti-CTLLA-4 and anti-PD1 affects amygdala-hippocampal circuit-dependent fear conditioning and memory consolidation process (5, 10) . The experimental setup includes a behavioral conditioning chamber (17.5 × 17.5 × 18 cm, Coulbourn Instruments) with steel shock floors (3.2 mm diameter slats, 8 mm spacing), a waste collection tray sprayed with 10% vinegar, dim light and a well established shock-tone system connected to the chamber (5, 10). During the first habituation day (Day 1), mice were allowed to habituate to the chamber for two minutes. Three pairs (evenly spaced at two-minute intervals) of auditory conditioned stimulus (CS; 16 kHz tone, 80 dB, lasting 120 sec) co-terminating with a mild foot shock unconditioned stimulus (US; 0.6 mA, 1 sec). Twenty-four hours later, on the subsequent 3 days of extinction training phase (Days 2-4), mice were initially habituated to the same contextual environment (dim light and vinegar odor) for two minutes before being presented with 20 non-US reinforced CS tones (16 kHz, 80 dB,lasting 120 sec, at 5 sec intervals). Twenty-four hours later (Day 5), fear testing was administered by presenting the mice with only three non-US reinforced CS tones (16 kHz, 80 dB, lasting 120 sec) at two minute intervals in the same context (hippocampal-dependent). At 72 hours after completion of hippocampal-depedent extinction test, an mPFC-dependent extinction test was administered using three non-US reinforced CS tones (16 kHz, 80 dB, lasting 120 sec) at two minute intervals in a differenet environment inclduing a white acrylic plate as a floor, new odor cue (10% almond in water), and an additioanl house light. Animals’ freezing behavior was recorded using a ceiling-mounted camera in the FE test chamber and recorded by an automated, video-based, motion measurement program (FreezeFrame, Coulbourn Instruments). FreezeFrame algorithms calculated a motion index for each frame of the video with higher values representing greater motion. To prevent biased analysis, an investigator blinded to the experimental groups set the motion index threshold representing immobility for each animal based on identifying a trough separating low values during immobility and higher values associated with motion. Freezing behavior was defined as continuous bouts of one second or more of immobility. The percentage of time each mouse spent freezing was then calculated for the final day of extinction test.

#### BrdU Preparation and Administration

One week after the last dose of the last immunotherapy injections, mice were administered with BrdU (5-Bromo-2′deoxyuridine, 50 mg/kg, I.P., once daily for 6 days, Sigma, Cat. B5002) to evaluate the impact of ICI treatment on *in vivo* neurogenesis. BrdU were pre-weighed into 75mg in six, carefully labeled, 50mL Falcon tubes (FisherSci, Cat. 14-432-22) and kept in -20°C until use. On the day of injection, one falcon tube was thawed to room temperature (RT) before processing. BrdU was dissolved in 30mL of 1X PBS (100mM, pH=7.6, FisherSci) and kept in a warm water bath (55-60°C) for 20 minutes. The solution was thoroughly vortexed until all visible particles were dissolved. Additionally, BrdU solution was filtered through Whatman® Puradisc 25 syringe filters (Sigma, Cat. WHA67502502) to remove any remaining undissolved BrdU particles. The solution was allowed to cool down to RT before injecting to the mice with the dosage described previously.

### Brain Tissues Collection for Immunohistochemistry

Mice were deeply anesthetized by inhaling 1.8% isoflurane v/v (Dechra) and euthanized via intracardiac perfusion by flushing ice-cold 1X PBS + 10U/ml heparin (Sigma, pH=4) in the left atrium until the venous outflow was clear. Perfusion with 1X PBS + 10U/ml heparin was quickly replaced with 4% PFA (Sigma) until the mouse body turned stiff and whole brains were immediately extracted and soaked in vials filled with 4% PFA (Sigma) overnight. Whole brains were switched to store in 1X PBS-0.05% Sodium Azide (Sigma-Aldrich Cat. S2002, pH 7.4). In order to avoid crystal formation during cryo-sectioning, whole brains were dehydrated by being submerged in a sucrose concentration gradient (10% to 30% w/v, Sigma, Cat. S7903, pH=7.4). When ready to be processed for cryo-setioning, whole brains were embedded in O.C.T compounds (VWR, Cat. 25608903) to support and stabilize the brains during sectioning. Each brain was sectioned at 30 µm thickness (coronal) and stored as floating tissues in 24-well plates that had 1X PBS-0.05% Sodium Azide (Sigma-Aldrich, pH 7.4) in a 4°C fridge for future use.

### Immunohistochemistry

Floating brain sections (N=4 brains/group, 2 sections/ brain) with visible hippocampi were selected for immunohistochemistry (IHC). Primary antibodies include Rat anti Mouse CD68 (1:500, Bio-Rad, Cat# MCA1957; RRID: AB_322219), Rabbit anti IBA1 (1:500, Wako, Cat# 019- 19741; RRID: AB_839504), Mouse anti NeuN (1:500, Millipore, Cat# MAB377, RRID:AB_2298772), Rabbit anti cFos (1:500, Abcam, Cat# ab190289, RRID:AB_2737414), Mouse anti GFAP (1:500, FisherSci, Cat# MA5-12023, RRID:AB_10984338), Rat anti MBP (1:500, Millipore, Cat# MAB386, RRID:AB_94975), Mouse anti Synaptophysin (1:1000, Sigma, S5768), Rat anti BrdU (1:150, Abcam, Cat# ab6326, RRID:AB_305426), Rabbit anti NeuN (1:500, Millipore, Cat# MABN140, RRID:AB_2571567) and Mouse anti PSD-95 (1:1000, FisherSci, Cat# MA1-045; RRID: AB_325399). Secondary antibodies include Goat anti Rat AF 647 (1:1000, Abcam, Cat# ab150159; RRID: AB_2566823), Goat anti Rabbit AF 488 (1:500, FisherSci, Cat# a11008; RRID: AB_143165), Donkey anti Mouse AF 488 (1:350, FisherSci, Cat# A-21202, RRID:AB_141607), Donkey anti Rabbit AF 568 (1:350, Abcam, Cat# ab175470, RRID:AB_2783823), Goat anti Mouse AF 568 (1:500, Abcam, Cat# ab175473, RRID:AB_2895153), Goat anti Rat AF 568 (1:1000, Abcam, Cat# ab175476, RRID:AB_2813739), and Goat anti Mouse AF 647 (1:1000, Abcam, Cat# ab150115; RRID: AB_2687948).

The staining procedure begins with washing sections 3 times in phosphate buffered saline (1X PBS, 100mM, pH 7.4, FisherSci) and if necessary, antigen retrieval using citrate buffer incubation (10mM, pH 6.0 with 0.05% Tween-20, Sigma) 70°C for 45 minutes. Tissues then incubated in borate buffer (pH 8.5, 100mM, Sigma, Cat. B0394) for 10 minutes followed by three 5-minute washes in 1X PBS. Tissues were then blocked in serum (10% normal goat or donkey, NGS or NDS) in PBS with 0.01% Triton X-100 (NDS, Jackson ImmunoResearch Labs Cat# 017- 000-121, RRID: AB_2337258; NGS, Jackson ImmunoResearch Labs Cat# 005-000-121, RRID:AB_2336990) Following an hour of incubation in the blocking solution, the tissues were incubated in primary antibodies for glial or neuronal components. Primary antibodies were prepared in 0.01% Triton X-100 (Sigma) 3% NGS or NDS in PBS and incubated for at least 12 hours at 4°C in a shaker incubator (Enviro-Genie Scientific). The tissues were then washed in 1X PBS for 3 times. They then incubated in the secondary antibodies for one hour. Finally, sections were counterstained with DAPI nuclear dye in PBS for 15 min (1 µmol/L, FisherSci, Cat# D1306, RRID:AB_2629482). Sections were again washed in PBS then mounted on superfrost slides (FisherSci) using Antifade mounting medium (VectaShield Cat# H-1000-10).

### Microscopy and 3D Algorithm-based Volumetric Quantification

Laser-scanning confocal microscope (Nikon Eclipse Ti2 AX) was used to image the immunostained brain sections at high resolution (2048p, 24 µm thick z-stacks with 0.5 µm per z-stack). Acquisition of confocal z stacks, deconvolution, and 3D algorithm-based volumetric quantification of immunofluorescent punta was carried out as described in detail previously (Markarian et al., 2021). 3D volume surfaces were created and quantified for each antigen of interest. Dual IHC stains such as CD68/IBA1, and co-localization volumetric between the 3D surfaces of the two markers were determined and automatedly quantified by Imaris (Oxford instruments). All images for each IHC stain were uniformly applied with the same parameters to obtain unbiased analyses.

### Induction of Experimental autoimmune encephalomyelitis (EAE)

Wild-type male 10-12 weeks C57BL/6J mice after receiving the combined ICI treatment as described above were induced with active or adoptive transfer experimental autoimmune encephalomyelitis (EAE). Active EAE was induced via SQ injection containing 200 μg MOG35–55 peptide (MEVGWYRSPFSRVVHLYRNGK) emulsified in Complete Freund′s Adjuvant, supplemented with *Mycobacterium tuberculosis* H37Ra (Hooke Laboratories Cat #EK-0111) as previously reported (6). Briefly, SQ injection was given at dosage of 200 μl emulsion at two sites over the flank region. Additionally, on the day of EAE induction and 48 hours post-induction, 200 ng pertussis toxin (List Biologic Laboratories, Cat. 181) was injected intraperitoneally to each mouse. To induce adoptive-transfer (AT) EAE, WT donor mice were first actively immunized with MOG35-55 peptide and a single dose of pertussis toxin as described above. 11 days later, cells were isolated from the inguinal draining lymph node (DLN) and spleen, and were activated *in vitro* with 20 𝜇g/mL MOG35-55 peptide in the presence of 20 ng/ mL of rm IL-12 (p70, containing both p40 and p35 subunits) and 10 𝜇g/mL anti-IFNγ (Clone XMG1.2, BioLegend, to protect MOG- responsive cells from activation-induced cell death) for 72 hr. ∼20 x 10^6^ donor cells were injected (i.p) into each WT recipient mice as described previously (Jairaman et al., 2021). Nutra-Gel wet food (Bio-Serve, Cat. S4798) was provided on the floor of the cage, along with long sipper bottles to facilitate feeding after the onset of EAE Mice were carefully monitored daily to assess for any development of clinical signs using the following scoring system: 0 - no signs, 0.5 - partially limp tail, 1 - limp tail, 1.5 - limp tail and hind leg inhibition, 2 - hind limb paresis, 2.5 - one hind limb paralysis, 3 both hind limb paralysis, 3.5 - hind limb paralysis and weakness in forelimbs, 4 - tetraplegia, and 5 - moribund or euthanized due to severe paralysis (scored >3.5 for 2 consecutive days).

## Statistical Analysis

Statistical analyses were performed using one-way ANOVA to confirm overall significance (GraphPad Prism, v8.0, RRID:SCR_002798). All data are expressed as the mean ± SEM. For comparisons between the vehicle and ICI-treated groups for the cancer volume, non-parametric, two-tailed unpaired *t* tests were performed with Holm-Sidak’s correction. For the analysis of ICI treatment in cancer and non-cancer groups, two-way ANOVA along with Tukey’s multiple comparisons test were performed. The Kruskal-Wallis test was used for the analysis of cytokine and IHC data. All analyses considered a value of *P* ≤ 0.05 to be statistically significant.

---

**utbl1.**

| REAGENTS or RESOURCES | SOURCE | IDENTIFIER |
| --- | --- | --- |
| **Antibodies** |  |  |
| eBioscience™ Fixable Viability Dye eFluor™ 780 | ThermoFisher | Cat# 65-0865-18<br>RRID: N/A |
| Alexa Fluor® 700 anti-mouse/rat/human CD27 Antibody | Biolegend | Cat# 124240<br>RRID:AB_2810383 |
| BD Horizon™ BV480 Rat Anti-Mouse I-A/I-E | BD Biosciences | Cat# 566086,<br>RRID:AB_2869739 |
| PE/Fire™ 700 anti-mouse NK-1.1 Antibody | Biolegend | Cat# 108774<br>RRID:AB_3083260 |
| PE anti-mouse/human CD3e Antibody | ThermoFisher | Cat# 12-0031-83<br>RRID:AB_465497 |
| PerCP Anti-Mouse CD8a (53-6.7) | Tonbo Biosciences | Cat# 67-0081,<br>RRID:AB_2621882 |
| Brilliant Violet 421(TM) anti-mouse CD19 | Biolegend | Cat# 115549,<br>RRID:AB_2563066 |
| Brilliant Violet 605(TM) anti-mouse/human CD11b | Biolegend | Cat# 101257,<br>RRID:AB_2565431 |
| Brilliant Violet 650(TM) anti-mouse CD45.2 | Biolegend | Cat# 109835,<br>RRID:AB_11203374 |
| FITC Anti-Mouse CD4 (GK1.5) | Tonbo Biosciences | Cat# 35-0041,<br>RRID:AB_2621665 |
| Brilliant Violet 570™ anti-mouse Ly-6C Antibody | Biolegend | Cat# 128030,<br>RRID:AB_2562617 |
| Rabbit anti IBA1 | FUJIFILM Wako | Cat# 019-19741; RRID:<br>AB_839504 |
| Rat anti Mouse CD68 | Bio-Rad | Cat# MCA1957; RRID:<br>AB_322219 |
| Mouse anti GFAP | FisherSci | Cat# MA5-12023,<br>RRID:AB_10984338 |
| Rat anti BrdU | Abcam | Cat# ab6326,<br>RRID:AB_305426 |
| Rabbit anti NeuN | Millipore | Cat# MABN140,<br>RRID:AB_2571567 |
| Rabbit anti cFos | Abcam | Cat# ab190289,<br>RRID:AB_2737414 |
| Mouse anti NeuN | Millipore | Cat# MAB377,<br>RRID:AB_2298772 |
| Mouse anti Synaptophysin | Sigma | Cat# S5768; RRID:<br>AB_477523 |
| Mouse anti PSD-95 | FisherSci | Cat# MA1-045; RRID:<br>AB_325399 |
| Rat anti Myelin Basic Protein | Millipore | Cat# MAB386,<br>RRID:AB_94975 |
| Donkey anti Rat AF 488 | FisherSci | Cat# A-21208,<br>RRID:AB_2535794 |
| Goat anti Rabbit AF 488 | FisherSci | Cat# a11008; RRID:<br>AB_143165 |
| Goat anti Mouse AF 568 | Abcam | Cat# ab175473,<br>RRID:AB_2895153 |
| Goat anti Rat AF 568 | Abcam | Cat# ab175476,<br>RRID:AB_2813739 |
| Goat anti Rat AF 647 | Abcam | Cat# ab150159; RRID:<br>AB_2566823 |
| Goat anti Mouse AF 647 | Abcam | Cat# ab150115; RRID: AB_2687948 |
| Donkey anti Mouse AF 488 | FisherSci | Cat# A-21202, RRID:AB_141607 |
| Donkey anti Rabbit AF 568 | Abcam | Cat# ab175470, RRID:AB_2783823 |
| Anti-PD1 | BioXCells | Cat# BP0273 RRID: AB_2687796 |
| Anti-CTLA-4 | BioXCells | Cat# BP0032 RRID: AB_1107598 |
| Armenian Hamster IgG | BioXCells | Cat# BP0091, RRID: AB_1107773 |
| Rat IgG2a | BioXCells | Cat# BP0089, RRID: AB_1107769 |
| 10X DAPI | FisherSci | Cat# D1306, RRID:AB_2629482 |
| **Biological Samples** |  |  |
| D4M-3A Melanoma Cells | Dr. F. Maragonni | UC Irvine |
| **Chemical, peptides, and recombinant proteins** |  |  |
| 5-Bromo-2'deoxyuridine (BrdU) | Sigma | Cat# B5002 |
| Gibco™ DMEM (1X) + GlutaMAX™-I | FisherSci | Cat# 10-569-010 |
| Gibco™ FBS | FisherSci | Cat# 10-082-147 |
| Gibco™ TrypLE™ Express Enzyme | FisherSci | Cat# 12-605-010 |
| Gibco™ Hibernate™-A Medium | FisherSci | Cat# A1247501 |
| Gibco™ DPBS (no magnesium, no calcium) | FisherSci | Cat# 14-190-144 |
| Gibco™ 10X DPBS (no magnesium, no calcium) | FisherSci | Cat# 14-200-075 |
| Nutra-Gel wet food | Bio-Serve | Cat# S4798 |
| MOG35–55 peptide (MEVGWYRSPFSRVVHLYRNGK) | PolyPeptide Laboratory | N/A |
| Freund's Adjuvant, Complete | Sigma | Cat# F5881 |
| Nonviable BD Difco™ Adjuvants (*Mycobacterium tuberculosis* H37Ra) | FisherSci | Cat# DF3114-33-8BD |
| Pertussis toxin | List Biologicals | Cat# 181 |
| Isospire™ (isoflurane) Inhalation Anesthetic | Dechra | N/A |
| Heparin sodium salt from porcine intestinal mucosa | Sigma | Cat# H3149 |
| Gibco™ AIM V™ Medium | FisherSci | Cat# 12055083 |
| Cytiva Percoll™ Centrifugation Media | FisherSci | Cat# 45001748 |
| Gibco™ HBSS | FisherSci | Cat# 14-175-079 |
| Paraformaldehyde | Sigma | Cat# 158127 |
| Sodium Azide | Sigma | Cat# S2002 |
| Sucrose | Sigma | Cat# S7903 |
| O.C.T compounds | VWR | Cat# 25608903 |
| Normal Goat Serum (NGS) | Jackson ImmunoResearch Labs | Cat# 005-000-121, RRID:AB_2336990 |
| Normal Donkey Serum (NDS) | Jackson ImmunoResearch Labs | Cat# 017-000-121, RRID: AB_2337258 |
| VECTASHIELD® Antifade Mounting Medium | VectaShield | Cat# H-1000-10 |
| **Software and Algorithms** |  |  |
| Noldus EthoVisionXT 17 Tracking Software | Noldus | N/A |
| FreezeFrame | Coulbourn Instruments | N/A |
| FlowJo | Tree Star | N/A |
| Prism | GraphPad | N/A |
| Imaris 10.0 | Oxford Instruments | N/A |
| **Others** |  |  |
| 31 Gauge insulin syringe | BD Sciences | Cat# 328411-1 |
| Precision Glide Needle Only 18 Gauge, 1 1/2" | BD Sciences | Cat# 305195 |
| Electronic Digital Caliper | VWR | Cat# 62379-531 |
| 50 mL Falcon tubes | FisherSci | Cat# 14-432-22 |
| 15 mL Falcon tubes | FisherSci | Cat# 339650 |
| Whatman® Puradisc 25 syringe filters | Sigma | Cat# WHA67502502 |
| Noldus EthoVisionXT 17 Video Capture Hardware System | Noldus | N/A |
| LSRFortessa | BD Bioscience | N/A |
| Scientific Industries SI-1200 Enviro-Genie | Enviro-Genie Scientific | N/A |
| Nikon Eclipse Ti2 AX | Nikon | N/A |

## REFERENCES

1. Acharya, M.M., J.E. Baulch, T. Lusardi, B.D. Allen, N.N. Chmielewski, A.A.D. Baddour, C.L. Limoli, and D. Boison. 2016a. Adenosine Kinase Inhibition Protects against Cranial Radiation- Induced Cognitive Dysfunction. Frontiers in molecular neuroscience 9:1–10.

2. Acharya, M.M., J.E. Baulch, T.A. Lusardi, B.D. Allen, N.N. Chmielewski, A.A. Baddour, C.L. Limoli, and D. Boison. 2016b. Adenosine Kinase Inhibition Protects against Cranial Radiation- Induced Cognitive Dysfunction. Frontiers in molecular neuroscience 9:42.

3. Acharya, M.M., L.A. Christie, M.L. Lan, E. Giedzinski, J.R. Fike, S. Rosi, and C.L. Limoli. 2011. Human neural stem cell transplantation ameliorates radiation-induced cognitive dysfunction. Cancer Res 71:4834–4845.

4. Acharya, M.M., K.N. Green, B.D. Allen, A.R. Najafi, A. Syage, H. Minasyan, M.T. Le, T. Kawashita, E. Giedzinski, V.K. Parihar, B.L. West, J.E. Baulch, and C.L. Limoli. 2016c. Elimination of microglia improves cognitive function following cranial irradiation. Scientific reports 6:31545.

5. Acharya, M.M., V. Martirosian, N.N. Chmielewski, N. Hanna, K.K. Tran, A.C. Liao, L.A. Christie, V.K. Parihar, and C.L. Limoli. 2015. Stem cell transplantation reverses chemotherapy-induced cognitive dysfunction. Cancer Res 75:676–686.

6. Allen, B.D., L.A. Apodaca, A.R. Syage, M. Markarian, A.A.D. Baddour, H. Minasyan, L. Alikhani, C. Lu, B.L. West, E. Giedzinski, J.E. Baulch, and M.M. Acharya. 2019. Attenuation of neuroinflammation reverses Adriamycin-induced cognitive impairments. Acta Neuropathol Commun 7:186.

7. Allen, B.D., A.R. Syage, M. Maroso, A.A.D. Baddour, V. Luong, H. Minasyan, E. Giedzinski, B.L. West, I. Soltesz, C.L. Limoli, J.E. Baulch, and M.M. Acharya. 2020. Mitigation of helium irradiation-induced brain injury by microglia depletion. Journal of neuroinflammation 17:159.

8. Barker, G.R., F. Bird, V. Alexander, and E.C. Warburton. 2007. Recognition memory for objects, place, and temporal order: a disconnection analysis of the role of the medial prefrontal cortex and perirhinal cortex. J Neurosci 27:2948–2957.

9. Barker, G.R., and E.C. Warburton. 2011. When is the hippocampus involved in recognition memory? J Neurosci 31:10721–10731.

10. Blansfield, J.A., K.E. Beck, K. Tran, J.C. Yang, M.S. Hughes, U.S. Kammula, R.E. Royal, S.L. Topalian, L.R. Haworth, C. Levy, S.A. Rosenberg, and R.M. Sherry. 2005. Cytotoxic T-lymphocyte- associated antigen-4 blockage can induce autoimmune hypophysitis in patients with metastatic melanoma and renal cancer. Journal of immunotherapy 28:593–598.

11. Buder-Bakhaya, K., and J.C. Hassel. 2018. Biomarkers for Clinical Benefit of Immune Checkpoint Inhibitor Treatment-A Review From the Melanoma Perspective and Beyond. Front Immunol 9:1474.

12. Cheung, Y.T., T. Ng, M. Shwe, H.K. Ho, K.M. Foo, M.T. Cham, J.A. Lee, G. Fan, Y.P. Tan, W.S. Yong, P. Madhukumar, S.K. Loo, S.F. Ang, M. Wong, W.Y. Chay, W.S. Ooi, R.A. Dent, Y.S. Yap, R. Ng, and A. Chan. 2015. Association of proinflammatory cytokines and chemotherapy- associated cognitive impairment in breast cancer patients: a multi-centered, prospective, cohort study. Annals of oncology : official journal of the European Society for Medical Oncology / ESMO 26:1446–1451.

13. Christie, L.A., M.M. Acharya, V.K. Parihar, A. Nguyen, V. Martirosian, and C.L. Limoli. 2012. Impaired cognitive function and hippocampal neurogenesis following cancer chemotherapy. Clinical cancer research : an official journal of the American Association for Cancer Research 18:1954–1965.

14. Curran, M.A., W. Montalvo, H. Yagita, and J.P. Allison. 2010. PD-1 and CTLA-4 combination blockade expands infiltrating T cells and reduces regulatory T and myeloid cells within B16 melanoma tumors. Proc Natl Acad Sci U S A 107:4275–4280.

15. Darnell, E.P., M.J. Mooradian, E.N. Baruch, M. Yilmaz, and K.L. Reynolds. 2020. Immune-Related Adverse Events (irAEs): Diagnosis, Management, and Clinical Pearls. Curr Oncol Rep 22:39.

16. Das, S., and D.B. Johnson. 2019. Immune-related adverse events and anti-tumor efficacy of immune checkpoint inhibitors. Journal for immunotherapy of cancer 7:306.

17. Duong, S.L., F.J. Barbiero, R.J. Nowak, and J.M. Baehring. 2021. Neurotoxicities associated with immune checkpoint inhibitor therapy. Journal of neuro-oncology 152:265–277.

18. Evans, G.J., and M.A. Cousin. 2005. Tyrosine phosphorylation of synaptophysin in synaptic vesicle recycling. Biochemical Society transactions 33:1350–1353.

19. Fleming, B., P. Edison, and L. Kenny. 2023. Cognitive impairment after cancer treatment: mechanisms, clinical characterization, and management. BMJ 380:e071726.

20. Glantz, L.A., J.H. Gilmore, R.M. Hamer, J.A. Lieberman, and L.F. Jarskog. 2007. Synaptophysin and postsynaptic density protein 95 in the human prefrontal cortex from mid-gestation into early adulthood. Neuroscience 149:582–591.

21. Goldberg, S.B., S.N. Gettinger, A. Mahajan, A.C. Chiang, R.S. Herbst, M. Sznol, A.J. Tsiouris, J. Cohen, A. Vortmeyer, L. Jilaveanu, J. Yu, U. Hegde, S. Speaker, M. Madura, A. Ralabate, A. Rivera, E. Rowen, H. Gerrish, X. Yao, V. Chiang, and H.M. Kluger. 2016. Pembrolizumab for patients with melanoma or non-small-cell lung cancer and untreated brain metastases: early analysis of a non-randomised, open-label, phase 2 trial. The Lancet. Oncology 17:976–983.

22. Hargadon, K.M., C.E. Johnson, and C.J. Williams. 2018. Immune checkpoint blockade therapy for cancer: An overview of FDA-approved immune checkpoint inhibitors. Int Immunopharmacol 62:29–39.

23. Jairaman, A., S. Othy, J.L. Dynes, A.V. Yeromin, A. Zavala, M.L. Greenberg, J.L. Nourse, J.R. Holt, S.M. Cahalan, F. Marangoni, I. Parker, M.M. Pathak, and M.D. Cahalan. 2021. Piezo1 channels restrain regulatory T cells but are dispensable for effector CD4(+) T cell responses. Sci Adv 7:

24. Kang, H., L.D. Sun, C.M. Atkins, T.R. Soderling, M.A. Wilson, and S. Tonegawa. 2001. An important role of neural activity-dependent CaMKIV signaling in the consolidation of long-term memory. Cell 106:771–783.

25. Keith, D., and A. El-Husseini. 2008. Excitation Control: Balancing PSD-95 Function at the Synapse. Frontiers in molecular neuroscience 1:4.

26. Kipnis, J. 2016. Multifaceted interactions between adaptive immunity and the central nervous system. Science 353:766–771.

27. Lee, J.B., H.R. Kim, and S.-J. Ha. 2022. Immune Checkpoint Inhibitors in 10 Years: Contribution of Basic Research and Clinical Application in Cancer Immunotherapy. Immune Netw 22:

28. Liddelow, S.A., K.A. Guttenplan, L.E. Clarke, F.C. Bennett, C.J. Bohlen, L. Schirmer, M.L. Bennett, A.E. Munch, W.S. Chung, T.C. Peterson, D.K. Wilton, A. Frouin, B.A. Napier, N. Panicker, M. Kumar, M.S. Buckwalter, D.H. Rowitch, V.L. Dawson, T.M. Dawson, B. Stevens, and B.A. Barres. 2017. Neurotoxic reactive astrocytes are induced by activated microglia. Nature 541:481–487.

29. Lo, J.A., M. Kawakubo, V.R. Juneja, M.Y. Su, T.H. Erlich, M.W. LaFleur, L.V. Kemeny, M. Rashid, M. Malehmir, S.A. Rabi, R. Raghavan, J. Allouche, G. Kasumova, D.T. Frederick, K.E. Pauken, Q.Y. Weng, M. Pereira da Silva, Y. Xu, A.A.J. van der Sande, W. Silkworth, E. Roider, E.P. Browne, D.J. Lieb, B. Wang, L.A. Garraway, C.J. Wu, K.T. Flaherty, C.E. Brinckerhoff, D.W. Mullins, D.J. Adams, N. Hacohen, M.P. Hoang, G.M. Boland, G.J. Freeman, A.H. Sharpe, D. Manstein, and D.E. Fisher. 2021. Epitope spreading toward wild-type melanocyte-lineage antigens rescues suboptimal immune checkpoint blockade responses. Sci Transl Med 13:

30. Loane, D.J., A. Kumar, B.A. Stoica, R. Cabatbat, and A.I. Faden. 2014. Progressive neurodegeneration after experimental brain trauma: association with chronic microglial activation. Journal of neuropathology and experimental neurology 73:14–29.

31. Markarian, M., R.P. Krattli, Jr., J.D. Baddour, L. Alikhani, E. Giedzinski, M.T. Usmani, A. Agrawal, J.E. Baulch, A.J. Tenner, and M.M. Acharya. 2021. Glia-Selective Deletion of Complement C1q Prevents Radiation-Induced Cognitive Deficits and Neuroinflammation. Cancer Res 81:1732–1744.

32. McGinnis, G.J., and J. Raber. 2017. CNS side effects of immune checkpoint inhibitors: preclinical models, genetics and multimodality therapy. Immunotherapy 9:929–941.

33. Montay-Gruel, P., M.M. Acharya, K. Petersson, L. Alikhani, C. Yakkala, B.D. Allen, J. Ollivier, B. Petit, P.G. Jorge, A.R. Syage, T.A. Nguyen, A.A.D. Baddour, C. Lu, P. Singh, R. Moeckli, F. Bochud, J.F. Germond, P. Froidevaux, C. Bailat, J. Bourhis, M.C. Vozenin, and C.L. Limoli. 2019. Long- term neurocognitive benefits of FLASH radiotherapy driven by reduced reactive oxygen species. Proc Natl Acad Sci U S A 116:10943–10951.

34. Myers, J.S., A.C. Parks, J.D. Mahnken, K.J. Young, H.B. Pathak, R.V. Puri, A. Unrein, P. Switzer, Y. Abdulateef, S. Sullivan, J.F. Walker, D. Streeter, and J.M. Burns. 2023. First-Line Immunotherapy with Check-Point Inhibitors: Prospective Assessment of Cognitive Function. Cancers (Basel) 15:

35. Oppegaard, K., C.S. Harris, J. Shin, S.M. Paul, B.A. Cooper, A. Chan, J.A. Anguera, J. Levine, Y. Conley, M. Hammer, C.A. Miaskowski, R.J. Chan, and K.M. Kober. 2021. Cancer-related cognitive impairment is associated with perturbations in inflammatory pathways. Cytokine 148:155653.

36. Othy, S., A. Jairaman, J.L. Dynes, T.X. Dong, C. Tune, A.V. Yeromin, A. Zavala, C. Akunwafo, F. Chen, I. Parker, and M.D. Cahalan. 2020. Regulatory T cells suppress Th17 cell Ca(2+) signaling in the spinal cord during murine autoimmune neuroinflammation. Proc Natl Acad Sci U S A 117:20088–20099.

37. Paolicelli, R.C., G. Bolasco, F. Pagani, L. Maggi, M. Scianni, P. Panzanelli, M. Giustetto, T.A. Ferreira, E. Guiducci, L. Dumas, D. Ragozzino, and C.T. Gross. 2011. Synaptic pruning by microglia is necessary for normal brain development. Science 333:1456–1458.

38. Paolicelli, R.C., A. Sierra, B. Stevens, M.E. Tremblay, A. Aguzzi, B. Ajami, I. Amit, E. Audinat, I. Bechmann, M. Bennett, F. Bennett, A. Bessis, K. Biber, S. Bilbo, M. Blurton-Jones, E. Boddeke, D. Brites, B. Brone, G.C. Brown, O. Butovsky, M.J. Carson, B. Castellano, M. Colonna, S.A. Cowley, C. Cunningham, D. Davalos, P.L. De Jager, B. de Strooper, A. Denes, B.J.L. Eggen, U. Eyo, E. Galea, S. Garel, F. Ginhoux, C.K. Glass, O. Gokce, D. Gomez-Nicola, B. Gonzalez, S. Gordon, M.B. Graeber, A.D. Greenhalgh, P. Gressens, M. Greter, D.H. Gutmann, C. Haass, M.T. Heneka, F.L. Heppner, S. Hong, D.A. Hume, S. Jung, H. Kettenmann, J. Kipnis, R. Koyama, G. Lemke, M. Lynch, A. Majewska, M. Malcangio, T. Malm, R. Mancuso, T. Masuda, M. Matteoli, B.W. McColl, V.E. Miron, A.V. Molofsky, M. Monje, E. Mracsko, A. Nadjar, J.J. Neher, U. Neniskyte, H. Neumann, M. Noda, B. Peng, F. Peri, V.H. Perry, P.G. Popovich, C. Pridans, J. Priller, M. Prinz, D. Ragozzino, R.M. Ransohoff, M.W. Salter, A. Schaefer, D.P. Schafer, M. Schwartz, M. Simons, C.J. Smith, W.J. Streit, T.L. Tay, L.H. Tsai, A. Verkhratsky, R. von Bernhardi, H. Wake, V. Wittamer, S.A. Wolf, L.J. Wu, and T. Wyss-Coray. 2022. Microglia states and nomenclature: A field at its crossroads. Neuron 110:3458–3483.

39. Parihar, V.K., J. Pasha, K.K. Tran, B.M. Craver, M.M. Acharya, and C.L. Limoli. 2014. Persistent changes in neuronal structure and synaptic plasticity caused by proton irradiation. Brain structure & function

40. Rogiers, A., C. Leys, J. De Cremer, G. Awada, A. Schembri, P. Theuns, M. De Ridder, and B. Neyns. 2020a. Health-related quality of life, emotional burden, and neurocognitive function in the first generation of metastatic melanoma survivors treated with pembrolizumab: a longitudinal pilot study. Supportive care in cancer : official journal of the Multinational Association of Supportive Care in Cancer 28:3267–3278.

41. Rogiers, A., C. Leys, J. Lauwyck, A. Schembri, G. Awada, J.K. Schwarze, J. De Cremer, P. Theuns, P. Maruff, M. De Ridder, J.L. Bernheim, and B. Neyns. 2020b. Neurocognitive Function, Psychosocial Outcome, and Health-Related Quality of Life of the First-Generation Metastatic Melanoma Survivors Treated with Ipilimumab. J Immunol Res 2020:2192480.

42. Rogiers, A., L. Willemot, L. McDonald, H. Van Campenhout, G. Berchem, C. Jacobs, N. Blockx, A. Rorive, and B. Neyns. 2023. Real-World Effectiveness, Safety, and Health-Related Quality of Life in Patients Receiving Adjuvant Nivolumab for Melanoma in Belgium and Luxembourg: Results of PRESERV MEL. Cancers (Basel) 15:

43. Salvador, A.F.M., and J. Kipnis. 2022. Immune response after central nervous system injury. Semin Immunol 59:101629.

44. Sznol, M., and I. Melero. 2021. Revisiting anti-CTLA-4 antibodies in combination with PD-1 blockade for cancer immunotherapy. Annals of oncology : official journal of the European Society for Medical Oncology / ESMO 32:295–297.

45. Tarhini, A. 2013. Immune-mediated adverse events associated with ipilimumab ctla-4 blockade therapy: the underlying mechanisms and clinical management. Scientifica (Cairo) 2013:857519.

46. Usmani, M.T., R.P. Krattli, Jr., S.M. El-Khatib, A.C.D. Le, S.M. Smith, J.E. Baulch, D.Q. Ng, M.M. Acharya, and A. Chan. 2023. BDNF Augmentation Using Riluzole Reverses Doxorubicin- Induced Decline in Cognitive Function and Neurogenesis. Neurotherapeutics 20:838–852.

47. Vinnakota, J.M., R.C. Adams, D. Athanassopoulos, D. Schmidt, F. Biavasco, A. Zahringer, D. Erny, M. Schwabenland, M. Langenbach, V. Wenger, H. Salie, J. Cook, O. Mossad, G. Andrieux, R. Dersch, S. Rauer, S. Duquesne, G. Monaco, P. Wolf, T. Blank, P. Hane, M. Greter, B. Becher, P. Henneke, D. Pfeifer, B.R. Blazar, J. Duyster, M. Boerries, N. Kohler, C.M. Chhatbar, B. Bengsch, M. Prinz, and R. Zeiser. 2024. Anti-PD-1 cancer immunotherapy induces central nervous system immune-related adverse events by microglia activation. Sci Transl Med 16:eadj9672.

48. Xiang, Z., J. Li, Z. Zhang, C. Cen, W. Chen, B. Jiang, Y. Meng, Y. Wang, B. Berglund, G. Zhai, and J. Wu. 2022. Comprehensive Evaluation of Anti-PD-1, Anti-PD-L1, Anti-CTLA-4 and Their Combined Immunotherapy in Clinical Trials: A Systematic Review and Meta-analysis. Frontiers in pharmacology 13:883655.

## References for Supplemental Materials and Methods

1. Acharya MM, Baulch JE, Klein PM, Baddour AAD, Apodaca LA, Kramar EA, et al. New Concerns for Neurocognitive Function during Deep Space Exposures to Chronic, Low Dose-Rate, Neutron Radiation. eNeuro 2019;6

2. Barker GR, Bird F, Alexander V, Warburton EC. Recognition memory for objects, place, and temporal order: a disconnection analysis of the role of the medial prefrontal cortex and perirhinal cortex. J Neurosci 2007;27:2948–57

3. Barker GR, Warburton EC. When is the hippocampus involved in recognition memory? J Neurosci 2011;31:10721–31

4. Bourin M, Hascoët M (2003) The mouse light/dark box test. Eur J Pharmacol 463:55–65.

5. Milad MR, Quirk GJ. Fear extinction as a model for translational neuroscience: ten years of progress. Annual review of psychology 2012;63:129–51

6. Othy S, Jairaman A, Dynes JL, Dong TX, Tune C, Yeromin AV, Zavala A, Akunwafo C, Chen F, Parker I, Cahalan MD. Regulatory T cells suppress Th17 cell Ca2+ signaling in the spinal cord during murine autoimmune neuroinflammation. Proc Natl Acad Sci U S A. 2020 Aug 18;117(33):20088–20099.

7. Parihar VK, Allen BD, Tran KK, Chmielewski NN, Craver BM, Martirosian V, Morganti JM, Rosi S, Vlkolinsky R, Acharya MM, Nelson GA, Allen AR, Limoli CL (2015c) Targeted overexpression of mitochondrial catalase prevents radiation-induced cognitive dysfunction. Antioxid Redox Signal 22:78–91.

8. Parihar VK, Maroso M, Syage A, Allen BD, Angulo MC, Soltesz I, Limoli CL (2018) Persistent nature of alterations in cognition and neuronal circuit excitability after exposure to simulated cosmic radiation in mice. Exp Neurol 305:44–55.

9. Petit-Demouliere B, Chenu F, Bourin M (2005) Forced swimming test in mice: a review of antidepressant activity. Psychopharmacology 177:245–255.

10. Schubert I, Ahlbrand R, Winter A, Vollmer L, Lewkowich I, Sah R. Enhanced fear and altered neuronal activation in forebrain limbic regions of CX3CR1-deficient mice. Brain Behav Immun 2018;68:34–43

