## Supplementary material for "Immune Checkpoint Inhibition-related Neuroinflammation Disrupts Cognitive Function": All_Supplemental_Figures

**Supplemental Figure S1**

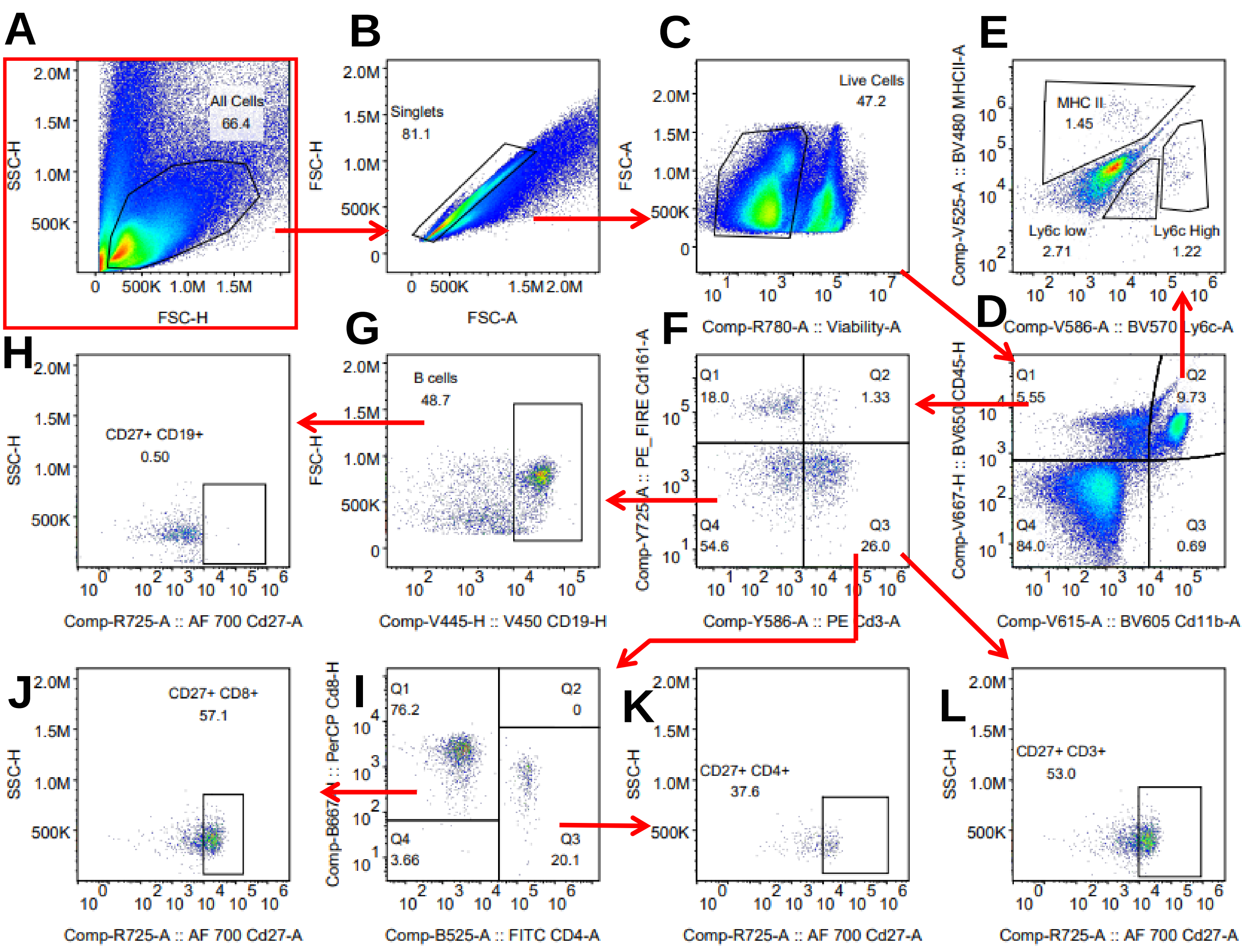


**Supplemental Figure S1. Gating strategy for immunophenotyping cells in the brain.** Sequential gating strategy to identify immune cells, exclude doublets, select live cells, and identify various immune cell subsets **(A-C)**. First, CD45 and CD11b markers **(D)** were used to gate lymphoid and myeloid populations. Q2 of **(D)** was used to further distinguish myeloid populations including myeloid cells expressing MHCII as well as Ly6c. Q1 of **(D)** was used for gating of NK, T, and NK T cells by plotting **(F)** CD161 and CD3. **(L)** All memory T cells were quantified by gating on cells from **(F)** Q3 that were CD27$^+$. **(I)** T cell subtypes were identified by plotting CD8 by CD4. **(J)** Memory CD8$^+$ T cells were identified by gating the CD27+ from **(I)** Q1. Memory CD4$^+$ T cells were identified by gating the CD27$^+$ from **(I)** Q3. Gate **(F)** -Q4 was used to identify **(G)** B cells. **(H)** Memory B cells were then identified by gating those cells that were CD27$^+$.

### Supplemental Figure S2

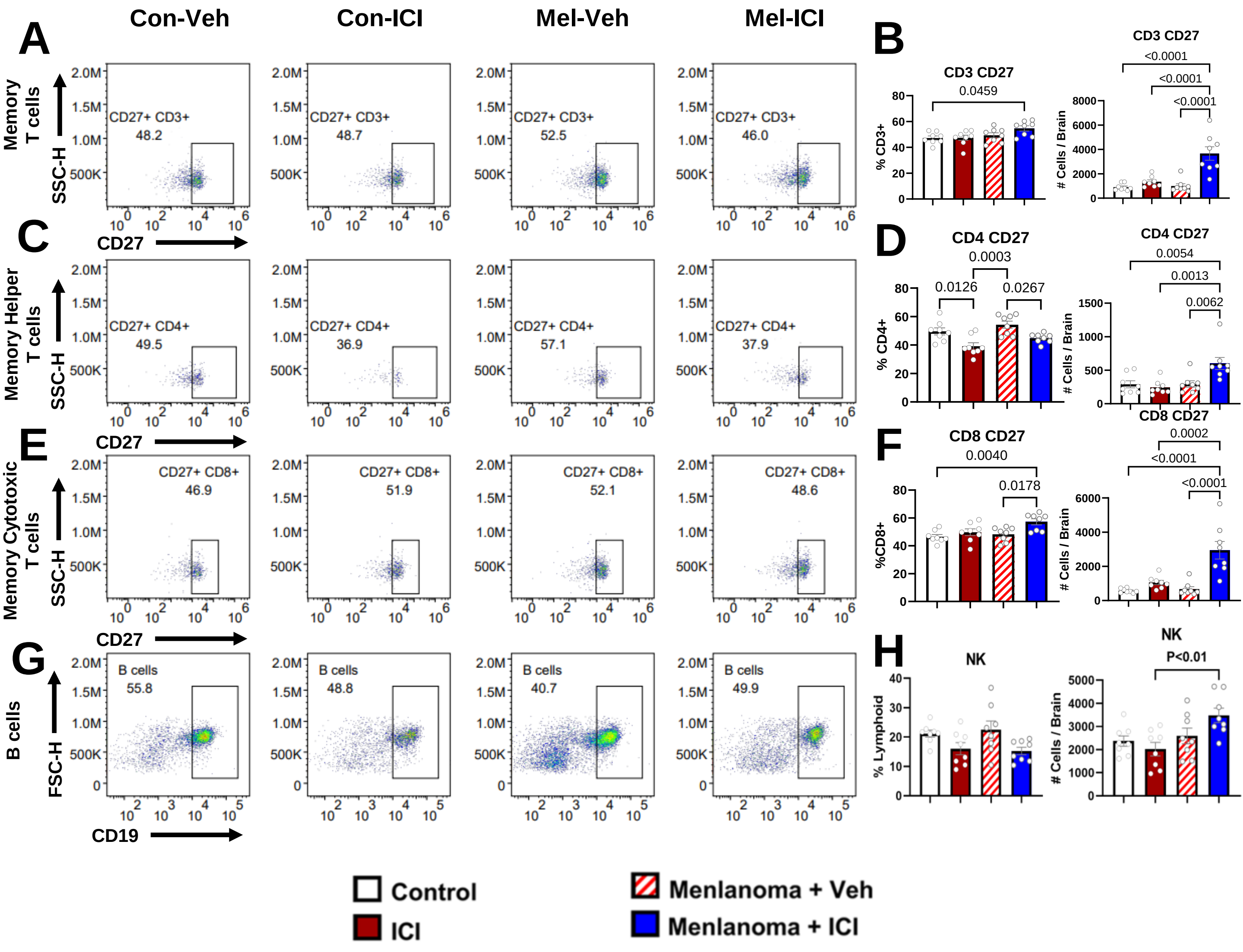


**Supplemental Figure S2. Immune cell profiling after CTLA-4 and PD-1 blockade.**

Animal experimental detail is described in **Fig. 1A**.

**(A)** Representative dot plots of $CD27^+$ $CD3^+$ T cell gating.

**(B)** Frequencies (% of $CD3^+$) and total number of $CD27^+$ $CD3^+$ T cells from **(A)**.

**(C)** Representative dot plots of $CD27^+$ $CD4^+$ T cell gating.

**(D)** Frequencies (% of $CD4^+$) and total number of $CD27^+$ $CD4^+$ T cells from **(C)**.

**(E)** Representative dot plots of $CD27^+$ $CD8^+$ T cell gating.

**(F)** Frequencies (% of $CD8^+$) and total number of $CD27^+$ $CD8^+$ T cells from **(E)**.

**(G)** Representative dot plots of $CD19^+$ B cell gating for graphs are shown in **Fig. 1H**.

**(H)** Frequencies (% of lymphoid) and total number of natural killer cells.

#### Supplemental Figure S3

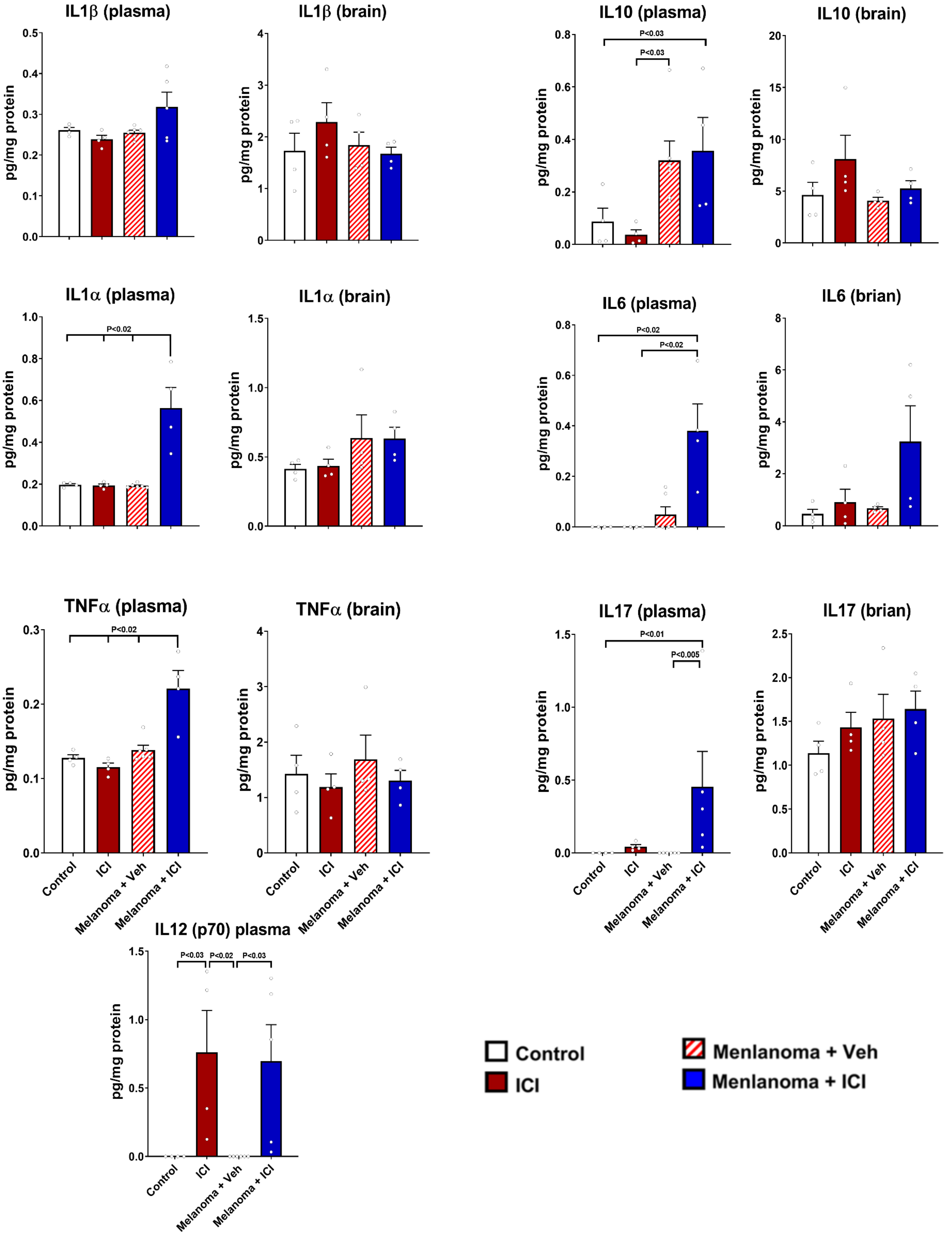

**Supplemental Figure S3. Effet of combined anti-CTLA-4 and anti-PD-1 on brain and plasma cytokines.**

Control and melanoma-bearing mice underwent 3 weeks of ICI treatment as described in previous experiments (Fig. 1). At 72 hours post-ICI treatment, animals were euthanized, blood (plasma) and brains (micro-dissected hippocampus) were collected for the cytokine ELISA. Brain tissue were homogenized in presence of protease inhibitor and centrifuged. Brain supernatants and plasma were assayed for cytokines using a magnetic bead-based customized kit (Thermo Fisher Scientific). Data is presented as mean ± SEM (N=4-6 mice/group). *P* values were derived from Kruskal-Wallis test

### Supplemental Figure S4

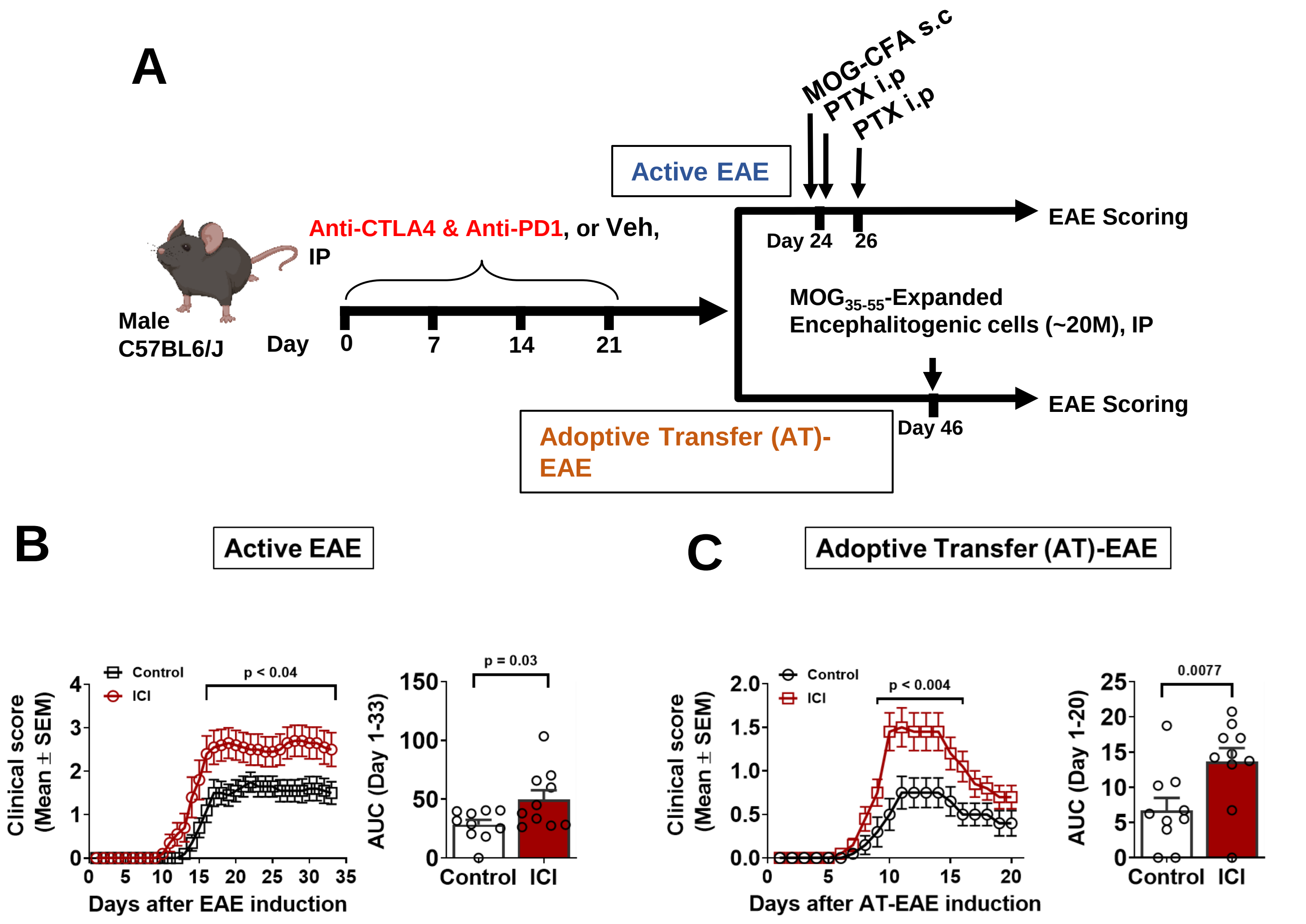


**Supplemental Figure S4.**

**ICI Predisposes CNS for Heightened Autoimmune Encephalomyelitis**

**(A)** Experimental Design: Mice underwent 3 weeks of ICI treatment as described in previous experiments (Fig. 1) followed by one month of recovery to allow ICI antibodies to clear. Animals then underwent active or passive (adoptive transfer) EAE induction. **(B)** Time-course plot of clinical scores following active EAE induction and the area under the curve (AUC). **(C)** Time-course plot of clinical scores following adoptive transfer EAE induction and the AUC.
